# Hybridization underlies localized trait evolution in cavefish

**DOI:** 10.1101/2021.05.11.443489

**Authors:** Rachel L. Moran, James B. Jaggard, Emma Y. Roback, Nicolas Rohner, Johanna E. Kowalko, C. Patricia Ornelas-García, Suzanne E. McGaugh, Alex C. Keene

## Abstract

Compared to selection on new mutations and standing genetic variation, the role of gene flow in generating adaptive genetic variation has been subject to much debate. Theory predicts that gene flow constrains adaptive evolution via natural selection by homogenizing allele frequencies among populations and introducing migrant alleles that may be locally maladaptive^1^. However, recent work has revealed that populations can diverge even when high levels of gene flow are present^2–4^ and that gene flow may play an underappreciated role in facilitating local adaptation by increasing the amount of genetic variation present for selection to act upon^5–8^. Here, we investigate how genetic variation introduced by gene flow contributes to adaptive evolution of complex traits using an emerging eco-evolutionary model system, the Mexican tetra (*Astyanax mexicanus*). The ancestral surface form of the Mexican tetra has repeatedly invaded and adapted to cave environments. The Chica cave is unique in that it contains several pool microenvironments inhabited by putative hybrids between surface and cave populations^9^, providing an opportunity to investigate the dynamics of complex trait evolution and gene flow on a local scale. Here we conduct high-resolution genomic mapping and analysis of eye morphology and pigmentation in fish from multiple pools within Chica cave. We demonstrate that hybridization between cave and surface populations contributes to highly localized variation in behavioral and morphological traits. Analysis of sleep and locomotor behaviors between individual pools within this cave revealed reduced sleep associated with an increase in ancestry derived from cave populations, suggesting pool-specific ecological differences may drive the highly-localized evolution of sleep and locomotor behaviors. Lastly, our analyses uncovered a compelling example of convergent evolution in a core circadian clock gene in multiple independent cavefish lineages and burrowing mammals, indicating a shared genetic mechanism underlying circadian disruption in subterranean vertebrates. Together, our results provide insight into the evolutionary mechanisms that promote adaptive genetic variation and the genetic basis of complex behavioral phenotypes involved in local adaptation.

## Main Text

A rapidly growing body of research demonstrates that gene flow, both among populations within species and between different species, is more common than previously thought and can play a key role in the evolutionary process by impeding or promoting adaptive divergence^5,6,10–12^. Hybrid zones resulting from interbreeding between lineages that occupy different environmental extremes offer a powerful means to detect targets of selection in the genome underlying complex, locally adapted traits. Natural variation present in recombinant hybrids can be leveraged through admixture mapping^13,14^, which identifies associations between genetic ancestry and trait variation in admixed populations formed by interbreeding between two or more diverged lineages. This association-based approach was first developed to uncover the genetic basis of diseases in humans following the observation that the frequencies of some disease-causing variants differ substantially among populations^15,16^. Recent advances in sequencing technology and statistical approaches have made it feasible to apply admixture mapping to identify adaptive loci underlying ecological divergence in plant and animal models of evolution^17–24^. However, previous studies in plants and animals have focused on hybrids formed between distinct species with substantial genetic divergence and reproductive isolation, making it difficult to identify regions associated with ecologically relevant traits versus intrinsic incompatibilities^25^. The application of admixture mapping to models of trait evolution has the potential to define fundamental interactions between genetic and environmental variation that shape evolution. Studies that apply whole genome sequencing to hybrids formed between interbreeding lineages in the earliest stages of divergence are likely to provide the most insight into the genetic basis of evolutionary change^24,26^, but are currently lacking.

The Mexican tetra, *Astyanax mexicanus*, is a powerful model system for investigating the genetic and evolutionary basis of trait development and behavior^27–31^. Surface populations inhabit rivers from Texas to Mexico and have invaded caves multiple times, resulting in at least 30 populations of cave-morphs in the Sierra de El Abra region of Northeast Mexico^9,32^. At least two independent lineages of surface fish, commonly referred to in the literature as “old” and “new” lineages, have invaded caves within the past roughly 200,000 years^33–37^. Cavefish populations have converged on numerous morphological traits that are thought to be adaptive in the cave environment, including albinism and eye loss^38^. In addition, cavefish have repeatedly evolved multiple behavioral changes, including sleep loss, which may increase time allocated to foraging in nutrient-poor cave environments^29,39^. Recently, the application of molecular genetic approaches has led to the identification of genetic factors that regulate some of these trait differences, but the evolutionary mechanisms underlying these genetic differences remain poorly understood.

Cave and surface populations are interfertile under laboratory conditions, and we recently identified a surprising amount of historical and contemporary gene flow between surface and cave populations^36^. The presence of admixture between populations raises the possibility that gene flow is a critical driver of trait evolution^36^. Unlike most caves, the Chica cave contains fish that appear to exhibit high levels of phenotypic variation, and fish are present across four pools that differ in proximity to the cave entrance, nutrient input, and physicochemical properties (e.g., dissolved oxygen) (Fig. 1A)^9,40–42^. Historical surveys suggested that fish exhibit a morphological gradation in troglobitic traits across pools, potentially shaped by environmental variation within the cave and ongoing influx of surface and cave morphs from underground waterways that feed into the cave^9^. Thus, this cave provides a natural system to study the effects of hybridization on trait evolution across a variable environment. Quantification of trait variation and formal tests for hybridization have yet to be conducted. Here we leverage robust differences in behavior and morphology between surface and cavefish populations of Mexican tetras, combined with whole genome sequencing, to investigate the ancestry of putative hybrids in Chica cave and to examine the genetic basis of trait variability across a heterogeneous environment.

**Figure 1.**
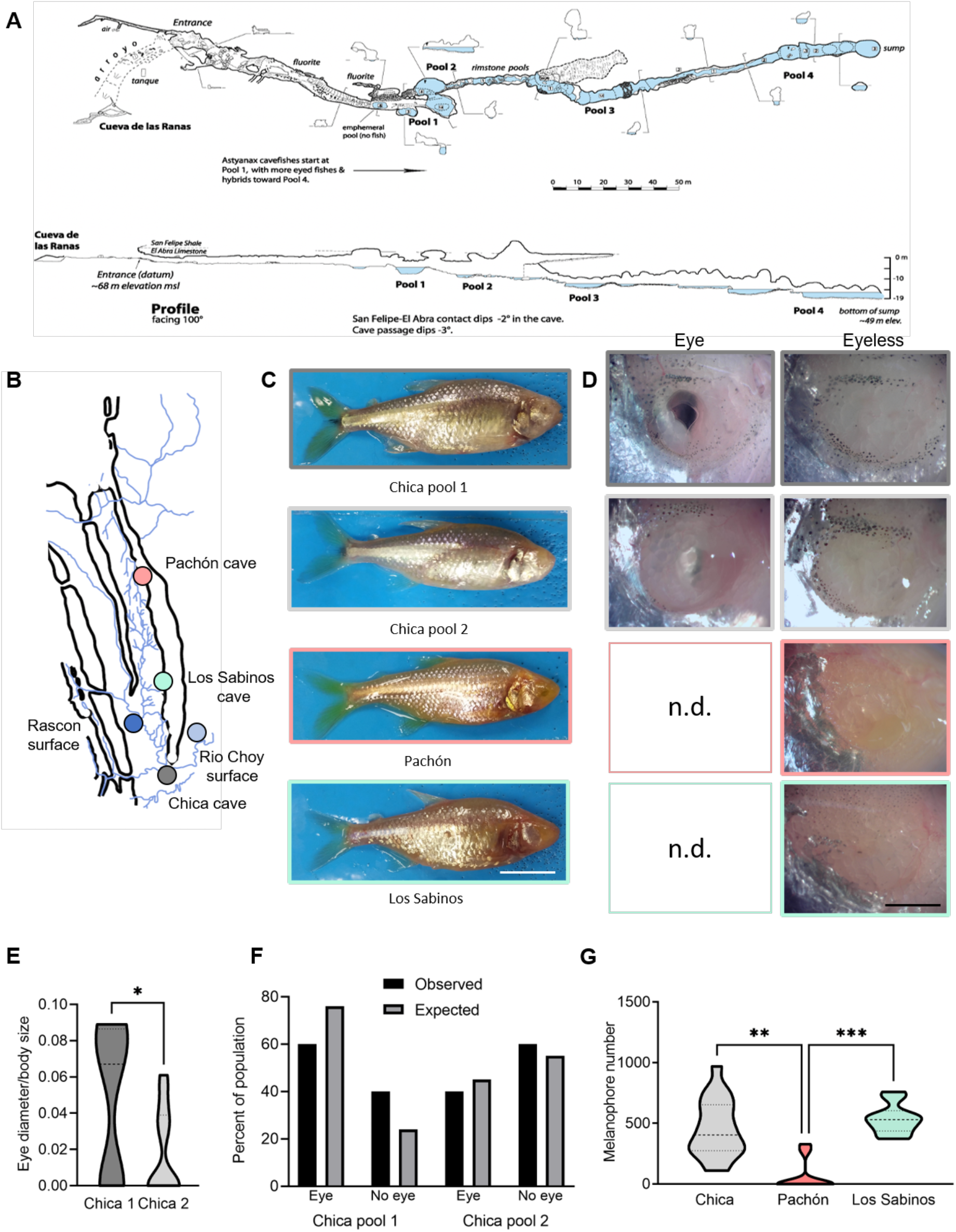
(A) Map of Chica cave modified with permission from ^43^. (B) Collection locations for cave and surface populations. For the two surface populations, the collection location for Río Choy is represented by a light blue circle and the collection location for Rascón is represented by a dark blue circle. (C) Representative images of wild-caught fish. Scale bar denotes 1 cm. (D) Representative images of eye morphology variations in Chica pools 1 and 2 and complete eye loss in wild-caught Pachón and Los Sabinos cave populations is denoted with “n.d.” for “no data” since there are no eyed fish present in these two populations. (E) Eye diameter is reduced in Chica pool 2 fish compared to pool 1 (*p <0.05, Unpaired t-test, t=1.88, df=17). Eye size was corrected to body length. (F) Eye morphology in Chica fish fits within expected outcome. Chica 1: observed 60% eye 40% no eye; expected 55% eye, 45% no eye, P>0.45 Binomial test. Chica 2: observed 40% eye, 60% no eye; expected 24% eye, 76% no eye, P>0.34 Binomial test. (D) Pigment quantification showing differences in melanin pigmentation in different populations (p <0.001, KW statistic=18.04, Kruskal-Wallis test with Dunn’s multiple comparison test: Chica vs Pachón, p < 0.001; Chica vs Los Sabinos, p=0.88; Pachón vs Los Sabinos, p < 0.001). Pigmentation are more variable among different cave populations, Brown-Forsythe test, P=0.03; Bartlett’s test, P=0.04.

We first conducted morphological and population genomic analysis to verify whether Chica fish represent hybrids between surface and cave populations. We also asked whether variation in morphological traits and allele frequencies are present between pool microenvironments within Chica cave. We collected adult fish from two adjacent pools in Chica cave that are partially hydrologically separate, referred to as Pool 1 and Pool 2 (Fig. 1A). We also collected adult fish from two non-admixed caves in the Sierra de El Abra regions, Pachón and Los Sabinos (Fig 1B,C). We scored wild-caught fish for two morphological traits, eye size and pigmentation, which previously have been qualitatively described as showing high variation within Chica cave compared to other cave populations. In agreement with laboratory stock populations, eyes were absent in wild-caught fish from Pachón and Los Sabinos caves (Figure 1D, Extended Data Figure 1). In contrast, the presence or absence of eyes was highly variable in wild-caught fish from both pools within Chica cave. Overall eye diameter was significantly larger in Chica Pool 1 fish compared to Pool 2 individuals (p <0.05, Unpaired t-test, t=1.69, df=17). Additionally, the sample from Chica Pool 1 contained more fish with eyes present (60%) than those with no eyes (40%) while the sample from Chica Pool 2 contained fewer fish with eyes (40%) and increased numbers with no eyes (60%) (Figure 1F). Binomial analysis demonstrated that the observed rate of the eye phenotype fit within expected outcome range (Chica Pool 1, p = 0.45; Chica Pool 2, p = 0.34).

We observed reduced melanin pigmentation levels in all cavefish, but these reductions vary among different cave populations. Pachón cavefish are considered largely albinic, while Los Sabinos retain vestigial melanocytes that result in a reduced pigmentation pattern compared to surface morphs^44,45^. Quantification of melanin pigmentation in wild-caught Pachón and Los Sabinos revealed comparable patterns to previous reports in lab-reared fish^44,45^ (Figure 1G). Although a number of pigmented individuals were present within the wild-caught Pachón and Los Sabinos populations, we observed low overall levels in the variability of melanin patterns within these cave populations. Interestingly, robust differences in the number of melnophores were observed between different populations of cavefish (Dunn’s multiple comparison, Chica vs.Pachon, p < 0.01; Pachon vs Los Sabinos, p < 0.01 Fig 1G; Extended Data Figure 1). Interestingly, the standard deviations between populations was significantly different, indicating that Chica cavefish are more variable in melanin morphology patterns (Brown-Forsythe test, p=0.03; Bartlett’s test, p=0.04). Taken together, these findings support the hypothesis that fish from Chica cave are the result of surface-cave hybridization and exhibit a high degree of phenotypic variability that differs between microenvironments within the Chica cave.

To identify the genetic basis of phenotype variability, we used whole-genome resequencing to conduct admixture analyses and genomic ancestry mapping in Chica cavefish to test for evidence of hybridization. We also used whole-genome resequencing to examine population structure between fish from Pools 1 and 2 within Chica cave, three other cave populations (Pachón, Los Sabinos, and Tinaja), and two surface populations (Río Choy and Rascón) (Fig. 2A,B). Our analyses revealed ongoing gene flow between Chica cave, Río Choy, and Tinaja cave populations (Fig. 2C), and confirmed that the Chica cave population represents a hybrid swarm resulting from over 2,000 generations of interbreeding between the nearby surface fish (from Río Choy/Tampaón) and southern El Abra cavefish (Fig. 2D; Extended Data Tables 7–8; Supplementary Information). This analysis indicated low levels of overall genome-wide divergence between fish from Chica cave Pool 1 and Pool 2, suggesting that gene flow is high among pools within Chica cave (Fig. 2E). Supporting this notion, all Chica individuals exhibited highly similar global ancestry proportions from surface versus cave parental populations (Pool 1 Cave Ancestry: Mean ± SE = 0.755 ± 0.004; Pool 2 Cave Ancestry: Mean ± SE = 0.756 ± 0.003; see Supplementary Information). Furthermore, the length distribution of ancestry tracts derived from the surface parental population did not differ between Chica pools (Extended Data Tables 7–8). Together, this indicates that gene flow from the surface population does not differ significantly between pools.

**Figure 2.**
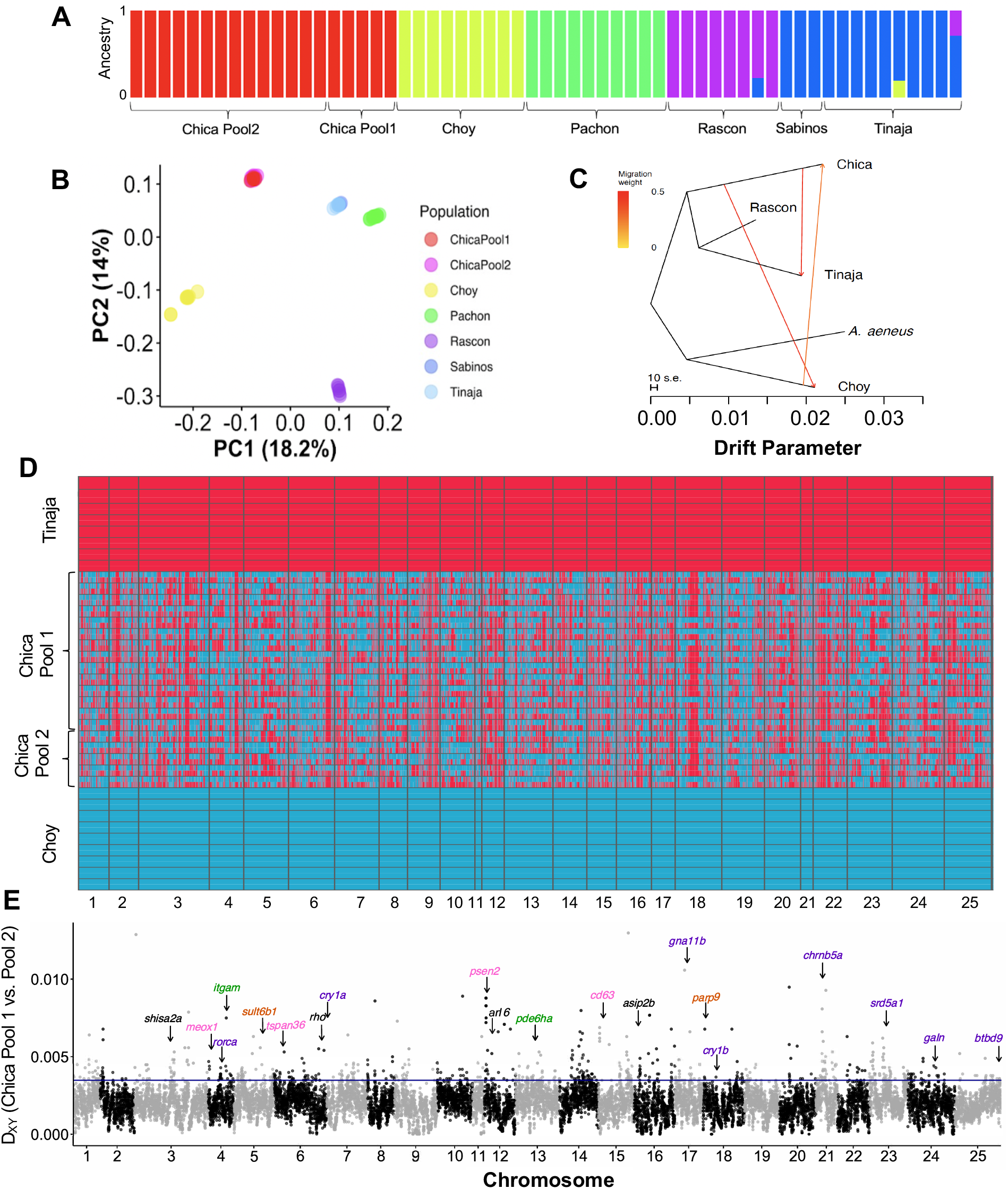
(A) ADMIXTURE barplot showing ancestry proportions for K=5. (B) Biplot of scores for the first two PCs. Note that individuals from Chica cave Pool 1 and Pool 2 overlap and individuals from Tinaja cave and Los Sabinos cave overlap. (C) Treemix tree with three migration events and rooted with the outgroup, *A. aeneus*. New lineage surface population (Río Choy) groups with *A. aeneus*, and old lineage surface (Rascón) and caves (Chica and Tinaja) all group together. Migration events are present between Chica cave and the geographically close surface population, Río Choy, and between Tinaja and Chica caves. (D) Local ancestry derived from surface (Río Choy, blue) versus cave (Tinaja, red) parental populations in hybrid fish from Chica cave. Each row represents a diploid individual with two haplotypes stacked on top of one another. (E) Absolute genetic divergence (Dxy) between fish from Chica cave Pool 1 versus Pool 2 in 50 kb windows across the genome. Locations are indicated for several top candidate genes with high divergence between Chica pools and biological functions related to sleep/circadian cycle (purple), eye size/morphology (green), metabolism (orange), and pigmentation (pink), or that are pleiotropically involved in two or more of these pathways (black) (see Extended Data Table 11). The 95^th^ percentile is delimited by a horizontal line.

Despite high levels of gene flow between pools, analysis of sequence divergence revealed several highly localized regions of genomic divergence between fish from adjacent pools within Chica cave, reflecting the morphological differences we observed between these pools (Fig. 2E). In genomic outlier windows where absolute genetic distance (i.e., Dxy) between Chica pools was in the top 5% of values, fish from Pool 2 were more likely to harbor alleles derived from the nearby southern El Abra cave populations (i.e., Tinaja) compared to Pool 1 (Wilcoxon rank sum test: W = 2.6511e+13, p < 2.2e-16). Specifically, 50.96% (371 out of 728) of the outlier windows with exceptionally high genetic divergence between pools contained a higher proportion of sites derived from cave (i.e., Tinaja) ancestry in Pool 2, whereas 39.56% (288 out of 728) had a higher proportion of sites derived from cave ancestry in Pool 1. The remaining 9.47% (69 out of 728) of outlier windows did not exhibit differences in ancestry between pools, which may be due to genetic differences that have accumulated via drift and/or regions that were not ancestry informative (i.e., the Hidden Markov Model was unable to accurately delimit ancestry blocks). We observed a positive correlation between the difference in local ancestry between pools and genetic distance between pools (Pearson’s correlation: r=0.0012, n=7,345,340, p=0.0011), indicating that the greater proportion of cave ancestry maintained in Pool 2 compared to Pool 1 drives genetic differences between the pools. Together, this demonstrates that gene flow has played a key role in driving genetic variation in this system and may facilitate evolution on a local scale.

Out of all functionally annotated protein-coding genes, those with a high degree of sequence divergence between pools (i.e., genetic distance in the top 5% of all genes) associated well with the characteristic suite of phenotypic differences typically present between non-admixed cave and surface populations (Extended Data Table 9). These genes with exceptional divergence between pools were significantly enriched for ontologies related to traits that differ between nona-dmixed surface and cave populations, including pigmentation (*bloc1s3, cd63, psen2, sox10, tspan36, meox1, asip2b, arl6, rab38c*), eye development and light detection (*c1qa, itgam, rho, pde6ha*), sensory processing by the lateral line neuromast (*rsph9, shisa2a*), metabolism (*parp9, sult6b1*), and sleep and the circadian cycle (*btbd9, srd5a1, mc3r, chrnb5a, galn, gna11b, rorca, cry1a, cry1b*) (Fisher’s exact tests, p < 0.05; Supplementary Information, Extended Data Tables 9–10). This set of 26 outlier genes with ontologies reatled to phenotypic differcne observed between non-admixed cave and surface populations provided strong candidates for local adaption. We used a deep convolutional neural network approach implemented in diploS/HIC^46^ to formally test for signatures of selection on these genes. We found that 21 (81%) of the 26 candidate genes occur within regions of the genome that appear to have experienced selective sweeps in one or both Chica pools (Extended Data Table 11). Furthermore, 20 (77%) of the 26 candidate genes for local adaptation also show extreme divergence (genetic distance in the top 5% of all genes) in one or more comparisons between cave and surface population pairs that do not show evidence of recent admixture (Extended Data Table 9), potentially pointing to general trends in the genetic underpinnings of cave evolution. These genes are strong candidates underlying the morphological differences in eye size and pigment we observed between pools within Chica cave, and suggest that adaptive behavioral (i.e., sleep) differences may also be present between pools. Taken together, our results indicate that hybridization may interact with varying selection pressures between different pool microenvironments within Chica cave to recapitulate phenotypic differences associated with divergent selection between cave and surface environments.

We observed a number of additional factors that further suggest the genetic differences identified in candidate genes for local adaptation in Chica cave could have functional consequences. We used *in silico* prediction with SIFT^47^ and VEP^48^ to identify mutations with deleterious effects that occur at higher frequencies in cavefish and re-analyzed transcriptional data obtained from Tinaja and Río Choy fry at 30 days post fertilization (dpf)^49^. These analyses revealed striking patterns of differential expression between lab-raised fry from surface versus cave populations and coding variants that affect protein function (see Supplementary Information, Extended Data Tables 11–13). Out of the 26 candidate genes with ontologies related to cave adapted phenotypes, 21 were expressed at 30 dpf. Of those 21 genes, 14 (67%) showed significant differential expression between cave and surface fish (Extended Data Table 13). We also identified putatively deleterious coding changes in five of our candidate genes with ontologies associated with sleep and the circadian cycle. One notable mutation is present in the gene cryptochrome circadian regulator 1a (*cry1a*), a transcriptional repressor. Cryptochromes play a highly conserved role in circadian clock regulation across plants and animals^50^. Knockout of *cry1a* results in defects in locomotor activity and behavioral rhythms in zebrafish^51^, and mutations in the human *cry1* ortholog are associated with a circadian rhythm sleep disorder (delayed sleep phase syndrome, DSPS)^52^. We observed that *cry1a* exhibits a nonsynonymous mutation, R263Q, that is present in Chica, Tinaja, and Pachón cave populations but not in Río Choy or Rascón surface populations. To determine whether this mutation is unique to the cavefish lineages, we examined an alignment of 284 CRY1 orthologs across 266 animal species (including invertebrates) downloaded from Ensembl (https://useast.ensembl.org/). We also downloaded the CRY1 ortholog for the Somalian cavefish *Phreatichthys andruzzi* that was available on NCBI (Accession: ADL62679.1). Remarkably, we found that the R263Q mutation is present in four distantly related cyprinid species from Somalia (*Phreatichthys andruzzi*)^53^ and China (the blind barbel *Sinocyclocheilus anshuiensis*, the golden-line barbel *Sinocyclocheilus grahami*, and the horned golden-line barbel *Sinocyclocheilus rhinocerous*), as well as two burrowing rodent species (the naked mole rat, *Heterocephalus glaber*, and the common degu, *Octodon degus*) (Fig. 3). *Phreatichthys and Sinocyclocheilus* cyprinid cavefish have convergently evolved troglomorphic traits that are shared by *Astyanax* characin cavefish, including reduction or loss of eyes and pigment, and disrupted circadian cycles^53–55^. The naked mole rat has also evolved many of the same characteristic traits associated with life in the dark, including reduced eye size and function, a disrupted circadian clock, and loss of sleep^56^. Our *in silico* analyses indicated that the R263Q mutation is predicted to be deleterious to protein function (Extended Data Table 11). This is supported by the observation that this position is otherwise highly conserved across plants and animals and occurs within the FAD binding domain of CRY^57,58^ (Fig. 3A). Our findings provide compelling evidence that the R263Q mutation in the core circadian clock gene *cry1* has convergently evolved up to five times in cavefish and burrowing mammals (Fig. 3B,C), indicating that a common genetic mechanism may contribute to disruption of sleep behavior and circadian rhythm in subterraneous vertebrates.

**Figure 3.**
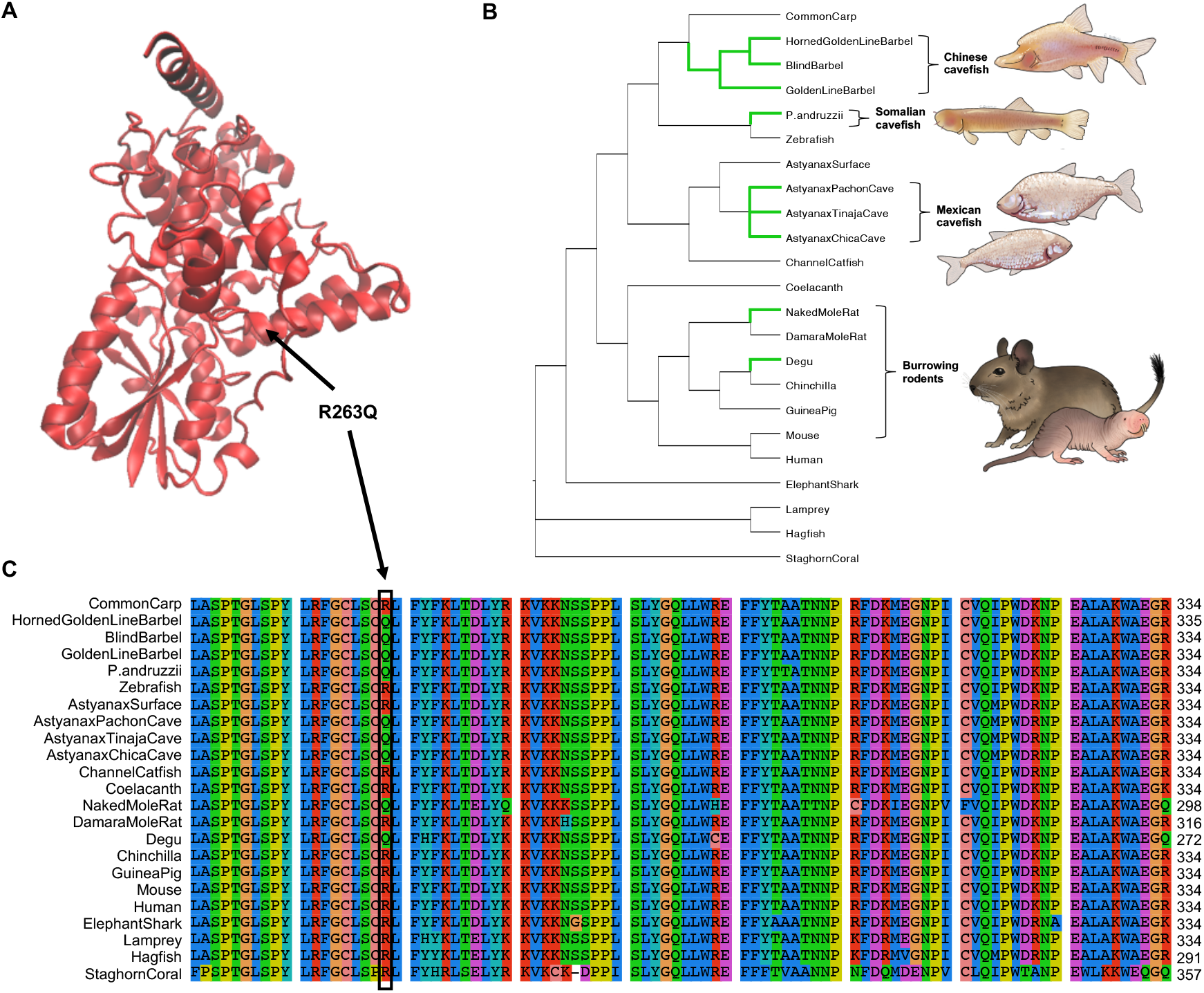
(A) Model of *Astyanax* Pachon cavefish CRY1A protein based on crystal structure of mouse CRY1 (PDB: 6kx7). The model for the *Astyanax* Pachon cavefish protein was generated with SWISS-MODEL and the comic structure was visualized with VMD (version 1.9.4). The location of R263Q (in the α10 within the FAD binding pocket) is indicated with an arrow. This image was made with VMD/NAMD/BioCoRE/JMV/other software support. VMD/NAMD/BioCoRE/JMV/ is developed with NIH support by the Theoretical and Computational Biophysics group at the Beckman Institute, University of Illinois at Urbana-Champaign. (B) Species tree for 23 animal species, selected to include subterranean lineages and their epigean relatives (based on the species tree available from Ensembl release 102 and ≡^5,59,60^). Branches where the R263Q mutation has evolved are highlighted in green. Illustrations depict *Astyanax* Mexican cavefish (Pachon cavefish top, Tinaja cavefish bottom), degu, and naked mole rat. (C) Section of multiple sequence alignment for CRY1 orthologs spanning sites 187 – 289 in the *Astyanax* CRY1A protein. The arginine to glutamine mutation at *Astyanax* site 263 is indicated with a black box.

Together, the enrichment of genes with ontologies related to sleep and the circadian cycle in our list of candidates with exceptionally high divergence between Chica pools and our observation of large-effect, cave-derived mutations in several of these genes suggests that sleep behavior may differ between fish from different pools within Chica cave. Multiple laboratory-bred cavefish populations exhibit convergence on sleep loss and increased locomotor activity^29,39^. While these behavioral differences are proposed to enhance foraging opportunity in nutrient-poor cave environments^28^, sleep has not been assayed in wild caught fish and it is not known whether sleep and activity differ based on local cave environments. To directly test whether the genomic differences we observed in candidate genes for sleep differences within Chica cave associate with functional differences, we analyzed behavioral variation in wild-caught fish from Chica Pool 1 and Pool 2. We also assayed non-admixed cavefish from Los Sabinos and Pachón for comparison. We measured sleep duration and locomotor activity in wild-caught fish under standard laboratory settings. We observed that total sleep in wild-caught Pachón cavefish is significantly reduced compared to Los Sabinos cavefish, similar to what is observed in fish derived from these populations in the laboratory (Fig. 4 A,B) (Dunn’s multiple comparison, p < 0.01). This provides evidence that the sleep loss observed in lab-reared stocks is replicated in wild-caught fish^39,61^. The duration of sleep in Chica fish from Pool 1 was significantly greater than sleep in fish from Pool 2 (Dunn’s multiple comparison, p < 0.05). The increased sleep duration from Chica Pool 1 fish was caused by an increase in number of sleep bouts compared to fish from both Chica Pool 2 and Pachón cave (Fig. 4B) (Dunn’s multiple comparison, Chica pool 2, p < 0.05, Pachon p < 0.05). Sleep bout length was reduced in fish from Chica Pool 2 and Pachón cave populations compared to fish from Chica Pool 1 (Fig 4C), though this comparison was not statistically significant. These differences in sleep cannot be explained by hyperactivity, as the average activity during periods of wakefulness (waking activity) did not differ between any of the populations (Fig. 4E). Therefore, hybrid Chica fish exhibit pool-specific differences in sleep, with fish from Pool 2 largely phenocopying Pachón cavefish and fish from Pool 1 exhibiting a greater sleep duration, similar to what has been previously observed in laboratory stocks of surface fish^29,39^. These results reveal the presence of behavioral differences between adjacent pools within Chica cave, with Pool 2 being more cavefish-like than Pool 1. This is in agreement with our genomic analyses, which found more cave ancestry maintained in Pool 2 compared to Pool 1 specifically in genomic regions of high divergence that contained genes related to phenotypes putatively implicated in local adaption in this system.

**Figure 4.**
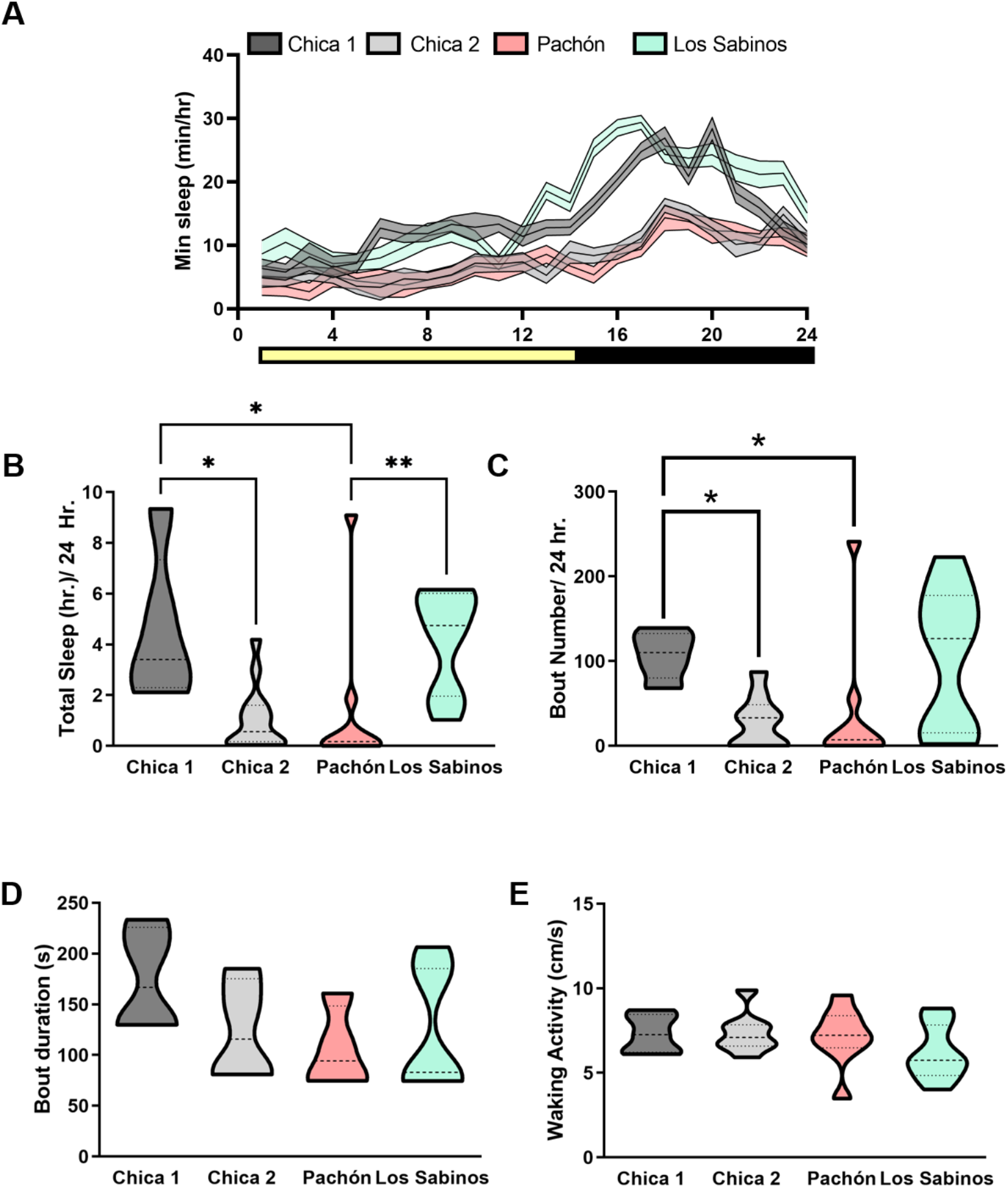
Sleep variation between and within wild-caught *A. mexicanus* cave populations. (A) Twenty-four hour sleep profiles in Chica Pool 1, Chica Pool 2, Pachón and Los Sabinos fish (B) Total sleep duration is variable among different populations of wild-caught fish (Kruskal-Wallis test, P < 0.001, KW statistic = 17.55). Chica Pool 1 fish sleep significantly longer than Chica Pool 2 fish (Dunn’s multiple comparison, p < 0.05). Wild-caught Pachón cavefish show reduced sleep compared to Chica Pool 1 (Dunn’s multiple comparison, p < 0.05). (C) Number of sleep bouts is variable in different cave populations (Kruskall-Wallis test, P < 0.05, KW statistic =10.62. Chica Pool 2 and Pachón fish have reduced sleep bout numbers compared to Chica Pool 1 (Dunn’s multiple comparison, Chica Pool 2, p < 0.05, Pachón, p < 0.05). (D) Sleep bout duration is not altered in any population of cavefish (Kruskal-Wallis test, P > 0.34, KW statistic = 3.46. (E) Waking activity is not altered among cave populations (Kruskal-Wallis test, P > 0.3, KW statistic = 3.65).

We conclude that genetic admixture across different environments produced substantially different phenotypes in very close geographic proximity. Our findings also imply that stronger selection against surface ancestry in Pool 2 of Chica cave could be maintaining differences in genes important to local cave pool via adaptation, rather than differential gene flow of each Chica pool and the local surface population. These results are remarkable given that the pools are separated by just 10 m. Additionally, we found that genes which are exceptionally divergent between the two cave pools associate well with the phenotypic differences observed. Notably, genes identified as highly divergent across pool microenvironments are also associated with phenotypes involved in nonadmixed surface and cave populations. These findings provide unprecedented insight into the genetic basis of local adaptation across varying environments in hybrid populations. Lastly, we identified a coding variant in the core circadian clock gene *cry1* that has convergently evolved in other distantly related cave-dwelling fish species and burrowing mammals. This suggests that a common genetic mechanism may contribute to disrupted circadian rhythms across multiple subterraneous vertebrate lineages that have adapted to live in constant darkness.

A rapidly growing body of work is demonstrating that introgressive hybridization often drives patterns of phenotypic evolution and may play an integral role in the evolutionary processes of local adaptation and speciation^12^. A number of studies have shown that behavioral variation can result from introgressive hybridization (e.g., song in hybrid Darwin’s finches^62^, mate choice in hybrid baboons^63^, defensive behavior in hybrid honey bees^23^), providing new substrate for selection to act upon. Hybridization has also been proposed to have influenced behavioral novelty in humans^64,65^. Here we mapped the genetic basis of complex behavioral differences resulting from the interplay between hybridization and selection in a vertebrate system. This demonstrates that hybridization may play an underappreciated role in shaping behavioral variation and evolution. Therefore, the identification of hybridization-mediated evolution in the *A. mexicanus* that inhabit Chica cave establishes this system as a model to study the genetic basis of evolution in complex behavioral and morphological traits.

## Methods

### Study system

Evidence suggests that there have been at least two colonization events of northern Mexico by surface dwelling *A. mexicanus*, typically referred to as the “old” and “new” lineage. One lineage of surface fish colonized the caves in the El Abra region and a separate lineage of surface fish subsequently colonized the northern Guatemala region and western Micos region caves of Northeastern Mexico. While we now know that these two lineages and their invasion of the caves were not timed in line with the “old” and “new” designations ^37^, we use this shorthand here since these labels are consistent with past work ^33,34,36,66^. The surface fish within the Rascón/Gallinas river system are most similar to the old lineage cavefish and were likely isolated from colonization by the new lineage surface fish due to a 105 m vertical waterfall ^37^. Cavefish within the El Abra region that descended from old lineage of surface ancestors are now within close geographic proximity to surface fish from the new lineage.

Fish occupy multiple pools within Chica cave that naturally differ in ecology. Whether the Chica cave population came from the old or new lineage stock has been the subject of much debate in the cavefish community. Fish from Chica cave show higher genetic differentiation from the rest of the El Abra cave populations, which some have interpreted as evidence of an independent invasion event ^34^. However, this pattern could also be explained by hybridization with local surface populations^67^. In accordance with this hypothesis, recent phylogenetic analyses have revealed that fish from Chica cave possess new lineage mitochondrial DNA and old lineage nuclear DNA, indicative of historical introgression ^33,35^.

Identifying the genetic underpinnings of behavioral evolution can be especially challenging in natural populations ^68,69^. A genomic signature of local adaptation is most detectable when gene flow is high among populations in different environments ^70,71^, as gene flow homogenizes the background level of divergence between populations while selection maintains differentiation at regions important to local adaptation. High levels of gene flow between the Chica cave and surface population together with strong selection for adaptation to the cave environment are predicted to shape patterns of divergence across the genome and provide insight into the genes important for maintaining cave phenotypes. Therefore, this system provides the unique opportunity to investigate the genetic basis of adaptive traits.

### Sequencing and Genotyping

We used whole genome resequencing and population genomic analyses to (1) characterize population structure and genetic relationships between and within the Chica cavefish, three other cavefish populations, and two surface populations (2) identify candidate regions for local adaptation with high levels of genetic differentiation between Chica pools, and (3) test for signatures of introgression between Chica cave and other nearby cave and surface populations. All sequencing used in these analyses originated from wild-caught fish collected from two adjacent pools within Chica cave (Pool 1, approximately 91 m from the entry, and Pool 2, approximately another 10 m into the cave; Fig. 1A).

Fin clips were collected from adult fish from Chica cave in 2015 and stored in 80% ethanol. We sequenced a total of 19 *A. mexicanus* samples from Chica cave (five from Pool 1 and 14 from Pool 2) using 125 bp paired end reads on an Illumina HiSeq 2500 at the University of Minnesota Genomics Center. Fin clips were collected from adult fish from Los Sabinos cave (n=3) in 2015 and were sequenced using 150 bp paired end reads on an Illumina NovaSeq S4. Genomic libraries for all Chica samples and two of the Los Sabinos samples (Sabinos_T3076_S26 and Sabinos_T3093_S27) were prepared using Illumina TruSeq v3 Nano DNA Sample Prep Kits. The genomic library for the third Los Sabinos sample (Sabinos1) was prepared using a Chromium Genome Library Kit and Gel Bead Kit v2 and a Chromium Genome Chip Kit v2. We obtained whole genome resequencing data for Pachón cavefish samples (n = 10) from a previously published study ^37^.

To investigate recent patterns of introgression between Chica cave and surface fish, we also obtained *A. mexicanus* sequence data from fish from one other cave population in the El Abra region that is not heavily admixed (Tinaja, n = 10), a nearby new lineage surface population (Río Choy, n = 9), and an old lineage surface population (Rascón, n = 8) ^37^. It has been hypothesized that caves within the southern El Abra region exchange migrants through subterranean connections, and Tinaja was previously shown to contain fish with mostly cave-like phenotypes. Thus, Tinaja cavefish sequence can provide a reference to identify cave alleles in the putative hybrid swarm present in Chica cave.

Río Choy contains new lineage surface fish and is a tributary of the Tampaón River, which is believed to be the source of surface fish in Chica cave. Rascón is a tributary of the Gallinas River, and contains old lineage surface fish^37^. Thus, including genomic data from Tinaja, Río Choy, and Rascón in our analyses provides a means to test for recent introgression between new lineage surface fish and old lineage cave fish within Chica cave. Previously published data from a closely related congener, *Astyanax aeneus* (n=1)^37^, was also included to serve as an outgroup in tests for introgression. Pachón, Tinaja, Río Choy, Rascón, and *A. aeneus* samples were all previously sequenced as 100 bp paired end reads on an Illumina HiSeq2000 at The University of Minnesota Genomics Center^37^. Raw sequencing data for these samples was downloaded from NCBI (SRA Accession Numbers SRP046999, SRR4044502, and SRR4044501).

We conducted genotype calling following the GATK Best Practices^72–74^(Extended Data Table 1). Adapters were trimmed from raw reads using Cutadapt v1.2.1. We trimmed samples for quality using Trimmomatic v0.30 and specified a minimum quality score of 30 across a 6 bp sliding window and discarded reads with a length of <40 nucleotides. Reads were aligned to the surface *Astyanax mexicanus* genome (Astyanax_mexicanus-2.0, downloaded from NCBI) using bwa v0.7.4^75^. We used Picard v2.3.0 (http://broadinstitute.github.io/picard/) to remove duplicates and add read group information and used samtools v1.7 ^76^ to split de-duplicated bams into mapped and unmapped reads. Mapped bams were used to generate per-individual gvcfs with the Genome Analysis Tool Kit (GATK) v3.7.0 HaplotypeCaller tool. We used the GenotypeGVCFs tool in GATK v3.8.0 to produce vcf files for each chromosome and unplaced scaffolds that include all individuals (and include invariant sites). The SelectVariants and VariantFiltration tools in GATK v3.8.0 were used to apply hard filters. We subset vcfs for each chromosome and unplaced scaffolds into invariant, SNPs, and mixed/indel sites and applied filters separately following GATK best practices (Supplementary Table 1). We then used the MergeVcfs tool in GATK v4.1.4 to recombine all subset VCFs for each chromosome and unplaced scaffold. Indels and the 3 bp region around each indel were removed using a custom python script. We used the vcftools^77^ --exclude-bed option to remove repetitive regions identified by WindowMasker and RepeatMasker^78^. We also used vcftools to only retain biallelic SNPs, to remove sites with greater than 20% missing data within each population, and to remove variants with a minor allele frequency <1%. This resulted in retaining a total of 225,462,242 sites throughout the genome, 3,337,738 of which were SNPs.

### Population Structure

To quantify the number of distinct genetic clusters (i.e., populations) present among the *A. mexicanus* cave and surface populations, we used ADMIXTURE v1.3.0 and Principal Components Analysis (PCA). For these analyses, we applied a more stringent missing data filter, only retaining sites with <10% missing data. To control for linkage between SNPs that cluster locally on a given chromosome, we thinned SNPs to 1 kb apart and did not include unplaced scaffolds. This resulted in a set of 678,637 SNPs. We ran ADMIXTURE for each value of K from two through nine and estimated the best value of K using the Cross Validation (CV) procedure in ADMIXTURE. The best K was chosen as the value that had the lowest CV error. We used Plink v1.90 to conduct the PCA. For this analysis, we again thinned SNPs to 1 kb apart, but included all placed and unplaced scaffolds. This resulted in a set of 733,979 SNPs.

We calculated absolute genetic divergence (Dxy) and relative genetic divergence (Fst) between populations and nucleotide diversity (Pi) within populations in non-overlapping 50kb windows across the genome using the python script popgenWindows.py (https://github.com/simonhmartin/genomics_general/blob/master/popgenWindows.py).

Fst can be influenced by heterogeneous genetic diversity between populations, and Herman et al.^37^ demonstrated that low Pi in caves can inflate relative divergence estimates in *A. mexicanus*. We therefore chose to use Dxy, which is not affected by levels of nucleotide diversity within populations, to identify regions of high genetic divergence between Chica pools.

We calculated Dxy on a site-by-site basis using a custom python script. This allowed us to calculate mean Dxy for each gene in the *A. mexicanus* genome annotation (v101, downloaded from ftp://ftp.ensembl.org/pub/release-101/gtf/astyanax_mexicanus/).

### Genome-wide tests for introgression

The population of fish within Chica cave has been hypothesized to be a hybrid swarm between cavefish originating from other caves in the El Abra region (which enter into Chica cave via a subterraneous connection) and surface fish from the nearby Río Choy/Tampaón river system^9^. To formally test this hypothesis, we conducted genome-wide tests for introgression between Chica cavefish and Tinaja cavefish and between Chica cavefish and Río Choy surface fish. We first used Treemix v1.13^79^ to confirm relationships between our focal populations and to visualize migration events between populations. Treemix builds a bifurcating tree to represent population splits and also incorporates migration events, which are represented as “edges” connecting population branches. We first built the maximum likelihood tree (zero migration events) in Treemix and then ran Treemix sequentially with one through five migration events. For this analysis, we included individuals from Chica, Río Choy (new lineage surface), Rascón (old lineage surface), and Tinaja (old lineage cave) *A. mexicanus* populations and the *A. aeneus* individual (outgroup), and SNPs were thinned to 1kb apart. We supplied this set of 700,502 biallelic SNPs to Treemix, rooted with *A. aeneus*, and estimated the covariance matrix between populations using blocks of 500 SNPs. Samples Tinaja_E, Tinaja_6, and Rascon_6 were excluded from this analysis because ADMIXTURE indicated that they were likely early generation hybrids. We calculated the variance explained by each model (zero through five migration events) using the R script treemixVarianceExplained.R^80^.

To test our hypothesis that Chica represents a hybrid population resulting from admixture between the nearby old lineage cave and new lineage surface populations, we used Dsuite v0.4^81^ to conduct formal tests for introgression between (1) Chica cavefish and Tinaja cavefish, and (2) between Chica cavefish and Río Choy surface fish. If no gene flow is occurring between the fish in Chica cave and the local surface population, we predict that fish from Chica (which has previously been shown to group phylogenetically with old lineage cavefish populations) should share more derived alleles with fish from Rascón (a surface population that is more geographically distant from Chica but also old lineage) than fish from Río Choy (a surface population that is geographically close to Chica cave but is new lineage). For this analysis, we supplied the set of 700,502 biallelic SNPs to Dsuite and specified *A. aeneus* as the outgroup. We again excluded three samples from Tinaja and Rascón with apparent hybrid ancestry. We used the Dsuite program Dtrios to calculate Patterson’s D statistic for all possible trios of populations using the ABBA-BABA test^82^. The ABBA-BABA test quantifies whether allele frequencies follow those expected between three lineages (e.g., sister species P1 and P2, and a third closely related species, P3) under expectations for incomplete lineage sorting (ILS). Observing a greater proportion of shared derived alleles between P1 and P3 but not P2 or between P2 and P3 but not P1 than what would be expected by chance (i.e., ILS) indicates introgression. Dsuite requires a fourth population, P4, to serve as an outgroup and determine which alleles are ancestral versus derived. Ancestral alleles are labeled as “A” and derived alleles are labeled as “B”. ABBA sites are those where P2 and P3 share a derived allele, and ABAB sites are those where P2 and P4 share a derived allele. The D statistic is calculated as the difference in the number of ABBA and BABA sites relative to the total number of sites examined. Dsuite uses jackknifing of the null hypothesis that no introgression has occurred (D statistic = 0) to calculate a p-value for each possible trio of populations.

Dsuite also calculates the admixture fraction, or f4-ratio, which represents the covariance of allele frequency differences between P1 and P2 and between P3 and P4. If no introgression has occurred since P1 and P2 split from P3 and P4, then f4 = 0. The f4 statistic is positive, this suggests a discordant tree topology indicative of introgression.

### Local Ancestry Inference

Hybrid genomes exhibit a mosaic of ancestry from their parental populations. A number of recent studies have shown that hybridization interacts with recombination and selection to shape patterns of local ancestry along chromosomes^83–87^. Non-random distributions of local ancestry in hybrid populations can indicate selection. Our goal here was to visualize patterns of introgression across the genome in Chica cavefish and determine whether more surface ancestry is present in Chica Pool 1 compared to Chica Pool 2. We used Hidden Markov Model (HMM) and fine-scale SNP mapping approaches to calculate ancestry proportions globally (i.e. genome-wide) and locally (at each base pair along each of the 25 chromosomes) in both Chica pools. To determine whether Pool 1 (nearer to the cave entrance) carries a higher proportion of surface ancestry compared to Pool 2 (deeper in the cave) at regions of the genome important to cave adaptation, we also asked whether regions of high divergence between pools exhibit higher differences in local ancestry.

We implemented a HMM-based approach in Loter ^88^ to infer genome-wide local ancestry in the Chica individuals. Tinaja and Río Choy served as the parental cave and surface populations, respectively, for the initial training stage of the HMM. We excluded two Tinaja samples that showed putative evidence of admixture^37^. This analysis allowed us to estimate global ancestry proportions and mean minor and major parent tract lengths for each individual. Ancestry tract lengths were converted from base pairs to Morgans using the median genome-wide recombination rate of median recombination rate of 1.16 cM/Mb obtained from a previously published genetic map for *A. mexicanus*^89^. We then estimated the number of generations since the onset of admixture (T_admix_) in each pool using the following equation:

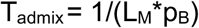

where L_M_ is the mean ancestry tract length from the minor parent in Morgans and p_B_ is the proportion of the genome derived from the major parent (the probability of recombining) ^90–92^.

We next used a chromosome painting approach with ancestry-informative sites to validate the delimitation of ancestry blocks detected by the HMM and to visualize patterns of introgression across the Chica cavefish genomes. This approach provides a lower level of resolution for ancestry block delimitation but with higher power to classify regions as derived from either parental genome. We identified alleles that were differentially fixed in Río Choy and Tinaja parental populations and had no missing data using the script get_fixed_site_gts.rb (https://github.com/mmatschiner/tutorials/blob/master/analysis_of_introgression_with_snp_data/src/get_fixed_site_gts.rb). We thinned SNPs to be a minimum of 1 kb apart and mapped these ancestry-informative sites in the Chica samples using the script plot_fixed_site_gts.rb (https://github.com/mmatschiner/tutorials/blob/master/analysis_of_introgression_with_snp_data/src/plot_fixed_site_gts.rb).

### Synthesizing patterns of genetic divergence and local ancestry

To quantify and visualize patterns of divergence between Pool 1 and Pool 2, we calculated summary statistics (Dxy, Fst, Pi) in non-overlapping 50kb windows across the genome using the python script popgenWindows.py (https://github.com/simonhmartin/genomics_general/blob/master/popgenWindows.py). We also calculated Dxy on a site-by-site basis using a custom python script (Cave_fish_Dxy.py). We asked whether there was an association between differences in local ancestry between pools and absolute genetic divergence (Dxy) between pools within outlier windows (which included coding and non-coding regions) and in coding regions alone.

We identified outlier windows as any 50kb window with a Dxy value above the 95th percentile (Dxy > 0.0035). Within each outlier window, we calculated the difference in local ancestry between fish from Pool 1 and Pool 2 at each site. We used a Wilcoxon rank sum test to identify whether ancestry differed within these regions between fish from Pool 1 versus Pool 2. We used Pearson’s correlation implemented in R (v4.0.2) to test for an association between difference in local ancestry and sequence divergence (Dxy) at each site between Chica Pool 1 and Pool 2 within outlier windows.

We calculated summary statistics for the coding region of each gene (i.e., max, median and mean Dxy, number of variant and invariant sites) within the *A. mexicanus* annotation (v101, downloaded from ftp://ftp.ensembl.org/pub/release-101/gtf/astyanax_mexicanus/) using a custom python script (Dxy_Summary_per_gene_ensemblGTF.py). This allowed us to rank genes by relative level of differentiation between Pool 1 and Pool 2. From this ranked list, we considered all genes with a mean Dxy above the 95th percentile (Dxy > 0.00276) as putative candidates for cave adaptation.

We used the GO Consortium Gene Ontology Enrichment Analysis tool (http://geneontology.org/) to ask whether any categories of biological processes were overrepresented in our set of outlier genes. We used the human (*Homo sapiens*) reference database for this analysis (20,851 genes). Fisher’s exact tests were performed to determine whether the number of genes associated with a given ontology were over- or under-represented in our set of outlier genes relative to the reference database.

We identified coding variants present among both Chica pools, Tinaja, and Río Choy and predicted the consequence of each variant on protein function using *in silico* computational analysis with the SIFT (sorting intolerant from tolerant) algorithm^47^ and the Ensembl Variant Effect Predictor (VEP) software suite^48^. SIFT uses sequence homology and data on the physical properties of a given protein to predict whether an amino acid substitution will be tolerated or deleterious. VEP performs annotation and analysis of genomic variants to predict impact on the protein sequence (i.e., modifier, low, moderate, or high).

Preliminary analyses indicated that one of our top candidate genes with high sequence divergence between Chica pools (*cry1a*) harbored a putative deleterious coding mutation (R263Q). To determine whether this variant is derived in cavefish and assess whether it occurs at evolutionarily conserved sites, we used the *Astyanax* surface fish genome annotation to obtain the CDS for *cry1a* from our population genomic data. We searched Ensembl for gene orthologs available in other animal species, including human, mouse, zebrafish, staghorn coral (*Acropora millepora*), thale cress (*Arabidopsis thaliana*), and three cyprinid cavefish species from China, the blind barbel (*Sinocyclocheilus anshuiensis*), the golden-line barbel (*Sinocyclocheilus grahami*), and the horned golden-line barbel (*Sinocyclocheilus rhinocerous*). We also downloaded the CDS for *cry1a* from another cyprinid cavefish species from Somalia, *Phreatichthys andruzzii*, from NCBI. We conducted a multiple species alignment for all 285 *cry1* orthologs using Muscle^93^. While investigating the R263Q mutation in *cry1a*, we identified a misassembly in the *Astyanax mexicanus* surface genome (Astyanax_mexicanus-2.0, downloaded from NCBI) affecting exons 9-13 of the *cry1a* coding region (*cry1a* CDS: 14,394-15,659 bp). Further investigation revealed that a portion of the coding region (*cry1a* CDS: 268-597 bp) was missing from the Pachon cavefish genome assembly (Astyanax_mexicanus-1.0.2, downloaded from NCBI). To confirm the mutation we identified in our population genomic data, we downloaded previously published *cry1a* mRNA sequences with complete CDS from Chica cave, Pachon cave, and Micos River (NCBI accession #s KF737846-KF737848). Aligning our population genomic data to the mRNA allowed us to verify that the correct exon coordinates were used around the mutation of interest. To visualize the location of the R263Q mutation, we created a 3D model of the *Astyanax* Pachon cavefish CRY1A protein in SWISS-MODEL^94^ using mouse CRY1 crystal structure (PDB: 6kx7). We imported the model into VMD (version 1.9.4) for visualization. To visualize the phylogenetic relationship between lineages with the R263Q mutation an identify putative instances of convergent evolution, we constructed a species tree that included 23 animal species (subterranean lineages and their close relatives) based on the species tree available from Ensembl release 102 and ^55,59,60^.

To test for signatures of selection in regions of the genome containing outlier genes, we used diploS/HIC^46^ to detect and classify selective sweeps. diploS/HIC uses a powerful supervised machine learning approach to identify windows in the genome that have undergone “soft” sweeps (selection on standing genetic variation) or “hard” sweeps (selection on new mutations) with high accuracy. We first simulated selective sweeps using discoal^95^ and then used the simulated data to train diploS/HIC. We provided diploS/HIC with a VCF containing the 3,337,738 SNPs showing <20% missing data across all populations and a masked version of the surface fish genome. We generated feature vectors for both Chica pools using the default settings of 11 sub-windows across a 1,100,000 Mb region (i.e., each window was 100,000 kb). diploS/HIC ran predictions using the feature vectors to classify each window as neutral (no evidence of a selective sweep), linkedSoft (loci near a window that has undergone a soft sweep), linkedHard (loci near a window that has undergone a hard sweep), Soft (loci that have undergone a soft sweep), or Hard (loci that have undergone a hard sweep). Windows lacking sufficient SNP data to make a prediction were labeled as “NA”.

To further investigate whether outlier genes between Chica pools are associated with phenotypic differences between cavefish and surface fish, we examined differential expression between lab-raised fry from Río Choy surface populations using a recently published RNAseq data set^49^ (SRA Project Accession #PRJNA421208). Briefly, batches of fry were sacrificed every 4 hrs between 6 am and 10 pm (i.e., 0 hrs, 4 hrs, 8 hrs, 16 hrs, 20 hrs; sample size mean ± SE across time points: n_Choy_ = 5.33 ± 0.33, n_Tinaja_ = 5.83 ± 0.17) at 30 days post-fertilization (dpf) for whole-body RNA extraction and sequencing^49^. Read counts for each gene across each sample were calculated as described in ^49^. We used DESeq2^96^ to calculate Log2(cavefish/surface fish) values. We considered genes to be differentially expressed if they had a Benjamini–Hochberg adjusted p-value < 0.05. Gene will less than 100 counts across all samples were excluded from the analyses.

### Fish collection and maintenance for phenotyping

We phenotyped wild-caught fish from Pools 1 and 2 within Chica cave and from two other cave populations, Pachón and Los Sabinos, which served as controls. Adult fish were collected from in 2015, during the dry season. The fish used in these analyses were the same fish that were sampled in 2015 for genomic sequencing analyses. The fish were transported and housed in the aquatic facility at Universidad Autónoma de Querétaro in 24 hour constant darkness. Fish were fed 1-2 times daily with dry flakes and kept at 23°C. These conditions were maintained throughout housing and experimental conditions for consistency. All fish were inspected for overall health, and any exhibiting signs of health or stress issues were excluded from experimental tests. All the samples were collected under the auspices of the permit SGPA/DGVS/0266/15, delivered by SEMARNAT. After the completion of behavioral assays, fin clips were collected from all Chica individuals for use in genomic sequencing as described above.

### Sleep behavior phenotyping

Fish were maintained in the lab for 8 months prior to behavioral assays. Adult fish were recorded in standard conditions in 10L tanks with custom-designed partitions that allowed for five fish (2L/fish) to be individually housed in each tank as previously described^39^. Recording chambers were illuminated with custom-designed IR LED source (Infrared 850 nm 5050 LED Strip Light, Environmental Lights). After a 4–5 day acclimation period, behavior was recorded for 24 hr beginning ZT0-ZT2. Videos were recorded at 15 frames/sec using a USB webcam (LifeCam Studio 1080 p HD Webcam, Microsoft) fitted with a zoom lens (Zoom 7000, Navitar). An IR high-pass filter (Edmund Optics Worldwide) was placed between the camera and the lens to block visible light. Videos were recorded using Virtualdub, a video-capturing software (Version 1.10.4) and were subsequently processed using Ethovision XT 9.0 (Noldus, IT). Water temperature and chemistry were monitored throughout recordings, and maintained at standard conditions in all cases. Ethovision tracking was set up as previously described^39^. Data was processed using Perl scripts (v5.22.0, developed on-site) and Excel macro (Microsoft)^39^. These data were used to calculate sleep information by finding bouts of immobility of 60 s and greater, which are highly correlated with increased arousal threshold, one of the hallmarks of sleep^39^.

### Morphological Characterization

Melanophores were quantified from bright-field images captured from each side of the body. Areas were chosen based on previous literature^97^ (i.e., caudal fin area, adipose fin area, dorsal area, eye cup area, anal fin area, infra-orbital area; see Extended Data Figure 1). Briefly, images were loaded into Fiji ImageJ (v. 1.7, National Insitutes of Health, Bethesda, MD). Images were color inverted in the selected area and using a preset noise tolerance allowed for melanophores to be automatically quantified by using pixel light intensity. If any melanophores were not counted, they were then manually added. Each image was analyzed by two different researchers to assure no significant discrepancies in quantifying, and the population of origin was blind to the researchers. All final quantifications were corrected to body length to account for different sized fish.

Eye presence and size were determined from images acquired on a handheld digital microscope (Dinoscope Pro AM4111T). Images were analyzed in Fiji ImageJ. Each image was inspected for the presence of an eye by two investigators, and the population of origin was blind to the researchers. Eye size was calculated in ImageJ by creating an ROI for the eye diameter and dividing this number by the length of the body to correct for overall size differences.

## Acknowledgements

We thank the University of Minnesota Genomics Center for their guidance and performing the cDNA library preparations and Illumina HiSeq 2500 sequencing. The Minnesota Supercomputing Institute (MSI) at the University of Minnesota provided resources that contributed to the research results reported within this paper. Funding was supported by NIH (1R01GM127872-01 to SEM, ACK, and NR) and NSF award IOS 165674 to ACK, and a US-Israel BSF award to ACK, and NSF IOS-1933076 to JK, SEM, and NR. Fish were collected under CONAPESCA permit PPF/DGOPA - 106 / 2013 to Claudia Patricia Ornelas García and SEMARNAT permit 02241 to Ernesto Maldonado. We thank the Mexican government for providing the collecting permit to R.B. in 2008 (DGOPA.00570.288108-0291).

## Supplementary Information

### Sequencing and Genotyping

Sequencing resulted in a mean ± SE of 187,777,319 ± 3,047,876 reads per individual for the 19 Chica samples and 331,445,356 ± 248,640,606 reads per individual for the three Los Sabinos samples. After quality filtering and mapping, all 60 samples had a mean ± SE genome-wide depth of coverage of 10.50 ± 0.53X (Extended Data Table 2).

### Population Structure

We examined population genetic structure among populations of cave and surface fish using PCA and ADMIXTURE. Populations examined (i.e., Río Choy, Los Sabinos, Tinaja, Rascón, Pachón, and Chica) generally showed separation from one another into distinct genetic clusters in both PCA and ADMIXTURE analyses (Figure 2A,B; Extended Data Figure 2). The first three principal components from the PCA explained over 45% of the total variance in the data (Extended Data Figures 2, 3; Extended Data Table 3), and separated the populations based on lineage and ecotype (surface or cave). For the ADMIXTURE analysis, comparison of cross validation error for K=2-9 indicated an optimal K of 5 (Extended Data Figure 4). Fish from Pool 1 and Pool 2 within Chica cave clustered together in the ADMIXTURE analysis and samples from both pools overlap completely in the PCA, indicating low overall levels of genetic divergence between the pools (Extended Data Figure 5). Los Sabinos and Tinaja are neighboring caves in the El Abra region and individuals from these clustered together in both analyses (Figure 2A,B; Extended Data Figure 2).

Average nucleotide diversity (Pi) across the genome did not differ between pools within Chica cave (Pi = 0.0021 for Chica Pool 1 and Pool 2; Extended Data Table 4). Notably, nucleotide diversity within Chica cave was 2-3X higher compared to other caves (Extended Data Table 4) and absolute genetic divergence between pools within Chica cave (Dxy = 0.0020) is comparable to that observed among other cave populations (Extended Data Table 5). Despite the fact that we observed no sites differentially fixed between Chica Pool1 and Pool 2, indicative that gene flow is ongoing, we observed several peaks of high sequence divergence between pools (Fig. 2E).

### Genome-wide tests for introgression

The results of two independent tests for introgression (implemented in Treemix and Dsuite) indicated hybridization between Chica cavefish and the nearby surface and cave lineages. Treemix first builds a bifurcating population tree and then fits for migration “edges” between branches. We chose the optimal number of migration events between populations in Treemix by examining the variance explained by models with zero through five migrations allowed. Adding one migration increased the proportion of variance explained from 0.59 to 0.92. The variance explained reached 1 and plateaued at three migration events (Extended Data Figure 6). Phylogenetic relationships were as expected based on previous analyses ^37^. Río Choy (new lineage) grouped with *A. aeneus*, and Rascón, Tinaja, and Chica (all old lineage) grouped together. We observed evidence of migration events between Río Choy surface fish with Chica cavefish and also between Tinaja cavefish and Chica cavefish (Figure 2C).

The results of the D statistic and f4 ratio tests implemented in Dsuite also confirmed introgression between Chica and Tinaja and between Chica and Río Choy. We were particularly interested in asking whether Chica has experienced recent gene flow with the local surface population, Río Choy. We observed that Río Choy and Chica share more derived sites (BBAA) than Chica and Rascón (ABBA) (Extended Data Table 6). Because Chica and Rascón populations are both derived from old lineage surface stock of *A. mexicanus*, whereas Río Choy is derived from new lineage surface stock, this pattern is indicative of introgression. Positive f4 ratios were also observed in all possible trios between Río Choy, Chica, Rascón, and Tinaja, indicative of introgression (Extended Data Table 6). Thus, our analyses support that Chica was originally colonized from an old lineage stock, but has subsequently experienced substantial introgression with new lineage surface populations.

### Local Ancestry Inference

To characterize patterns of introgression in Chica cavefish genomes, we used HMM-based ancestry inference and SNP mapping at ancestry-informative sites. The HMM-based approach implemented in Loter revealed that the genomes of individuals from Chica cave were composed of approximately 75% cavefish (i.e. Tinaja) ancestry and 25% surface fish (i.e. Río Choy) ancestry (Extended Data Figure 6). The two pools within Chica cave exhibited nearly identical global ancestry proportions corresponding to surface (Pool 1 Surface Ancestry: Mean ± SE = 0.245±0.004; Pool 2 Surface Ancestry: Mean ± SE = 0.244±0.003) versus cave (Pool 1 Cave Ancestry: Mean ± SE = 0.755±0.004; Pool 2 Cave Ancestry: Mean ± SE = 0.756±0.003) parental populations. Surface ancestry tract length distribution was similar in both Chica pools (Extended Data Table 7). The timing since the onset of admixture also did not differ between pools (Mean ± SE generations since hybridization: Pool 1 = 2315±19, Pool 2 = 2294±17) (Extended Data Table 8).

We identified 89,810 sites with complete fixation of different alleles in the parental populations (i.e., Río Choy and Tinaja). We mapped these sites across all 25 diploid chromosomes for each of the 19 Chica fish. In general, ancestry was highly admixed across Chica genomes and showed a pattern in agreement with the findings of the HMM-based ancestry inference (Fig. 2D; Extended Data Figure 7).

### Gene Expression

To investigate whether outlier genes may play a functional role in generating phenotypic variation associated with local adaptation, we examined differences in gene expression between lab-reared fry from Río Choy surface and Tinaja cave populations at multiple time points^49^. Out of a total set of 27,420 genes, 18,958 were expressed in fry at 30 dpf, and 11,901 (63%) of expressed genes showed significant differential expression (Benjamini–Hochberg adjusted p-value < 0.05) between cave and surface fish during at least one time point (Extended Data Tables 12–13). For the subset of 706 genes in the top 5 % of Dxy values between Chica Pool 1 and Pool 2, 586 were expressed in fry at 30 dpf, and 389 (66%) of expressed genes showed significant differential expression between cave and surface fish during at least one time point (Extended Data Tables 12–13). This represents an enrichment of differentially expressed genes in the outlier genes compared to the entire genome-wide data set (χ^2^ square test: χ^2^ _1,19,544_= 3.168, p = 0.031). Out of 26 top Dxy genes with ontologies related to cave adapted phenotypes (i.e., sleep/circadian cycle, light detection, eye size/morphology, metabolism, pigmentation), 21 were expressed at 30 dpf. Of those 21 genes, 14 (67%) showed significant differential expression between cave and surface fish during at least one time point (Extended Data Table 13).

### Phenotyping

The pattern we documented of more surface-like phenotypes near the entrance of the cave (in Pool 1) and more cave-like phenotypes deeper in the cave (in Pool 2) (Fig. 1C,D; Fig. 4) contrasts the findings of previous studies. Over eight decades ago,^42^ and ^41^_documented more cave-like phenotypes in the front of the cave (Pool 1) and more surface-like fish deeper in the cave (Pools 2-4). It was speculated that both cavefish and surface fish from nearby populations were entering Chica cave subterraneously near the lower pools, and that the connection between Pool 1 and Pools 2-4 was dynamic. A lack of connectivity between Pool 1 and the other pools was hypothesized to create a harsher, low nutrient environment towards the front of the cave, resulting in increased survival of fish with cave adapted phenotypes in this pool. However,^43^ recently suggested that surface fish may have access to the entrance of the cave via runoff from nearby drainages. Indeed, river-dwelling fish species (i.e., cichlids and poecilids) were observed near the entrance of the cave in Pool 1 when fish were collected for the present study in 2015. This suggests that surface fish may have access to Chica cave Pool 1 directly via the entrance and also via a deeper, subterraneous connection. Future studies using mark-recapture would provide a more definitive answer.

## Extended Data Tables

**Extended Data Table 1.** GATK filters applied to variant and invariant sites.

| Invariant sites | SNPs | Mixed/indels |
| --- | --- | --- |
| QD < 2.0 | QD < 2.0 | QD < 2.0 |
| FS > 60.0 | FS > 200.0 | FS > 200.0 |
| MQ < 40.0 | ReadPosRankSum < -20.0 | ReadPosRankSum < -20.0 |

**Extended Data Table 2.** Read counts and coverage info.

https://docs.google.com/spreadsheets/d/1_3eFqa2BuOZWyVwU9J1_vDAEt9D3s4ySBtOFSUow3Ek/edit#gid=0

**Extended Data Table 3.** Eigenvalues for PC 1-20 from PCA containing all six *Astyanax* populations (i.e., Chica, Río Choy, Los Sabinos, Pachón, Rascón, and Tinaja).

| PC | EigenValue |
| --- | --- |
| 1 | 3.49399 |
| 2 | 2.69367 |
| 3 | 2.48907 |
| 4 | 1.45645 |
| 5 | 1.17887 |
| 6 | 0.800319 |
| 7 | 0.796332 |
| 8 | 0.795117 |
| 9 | 0.789957 |
| 10 | 0.786389 |
| 11 | 0.764248 |
| 12 | 0.393822 |
| 13 | 0.390711 |
| 14 | 0.383791 |
| 15 | 0.378667 |
| 16 | 0.376847 |
| 17 | 0.337861 |
| 18 | 0.318567 |
| 19 | 0.2882 |
| 20 | 0.279671 |

**Extended Data Table 4.** Nucleotide diversity (Pi) within populations.

| Population | n | $P_i$ |
| --- | --- | --- |
| Chica Pool 1 | 5 | 0.0021 |
| Chica Pool 2 | 14 | 0.0021 |
| Río Choy | 9 | 0.0028 |
| Pachón | 10 | 0.0008 |
| Rascón | 10 | 0.0013 |
| Los Sabinos | 3 | 0.0007 |
| Tinaja | 8 | 0.0008 |

**Extended Data Table 5.**
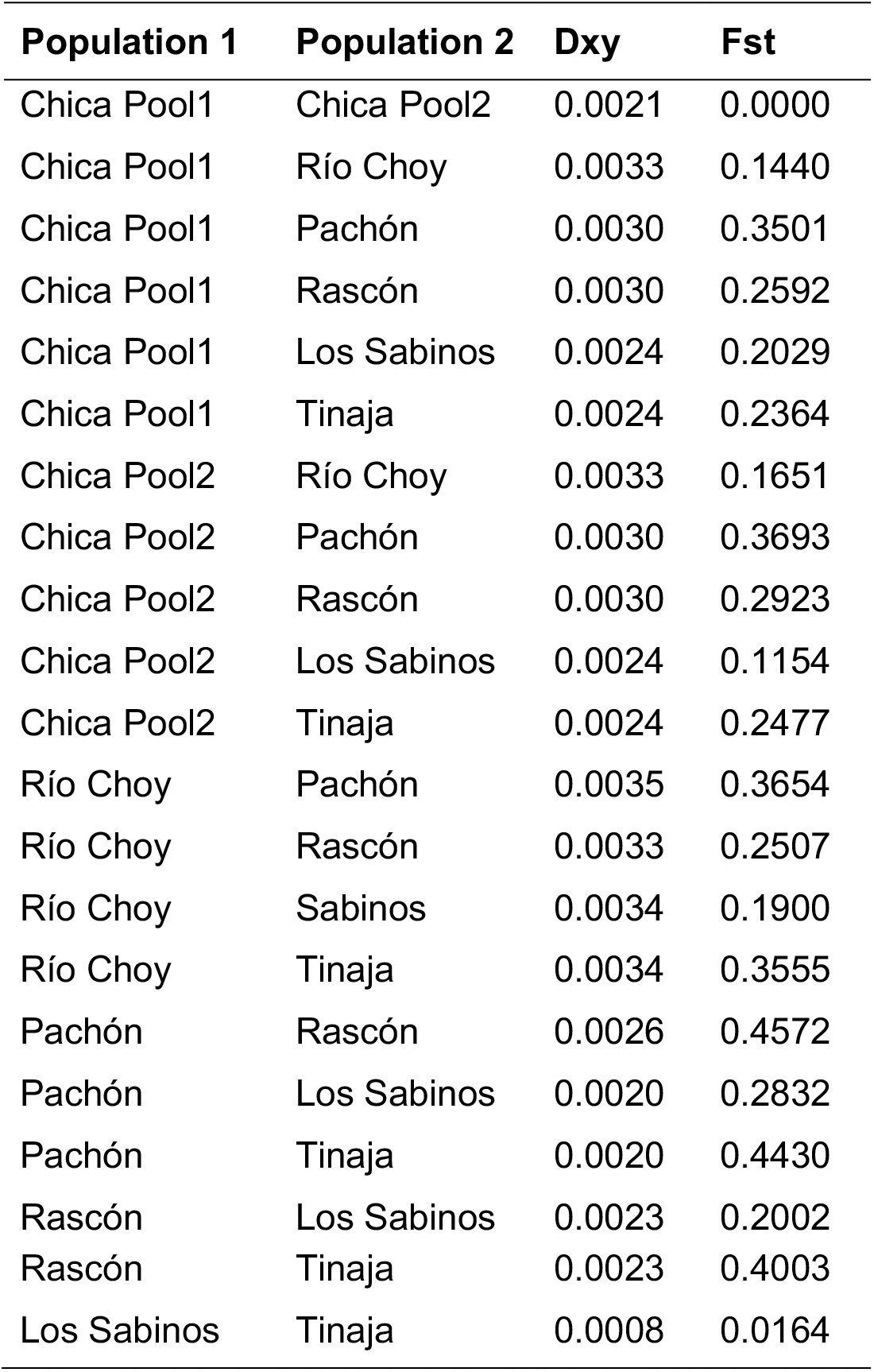
Absolute genetic divergence (Dxy) and relative genetic divergence (Fst) between populations. See Extended Data Table 4 for population sample sizes.

| Population 1 | Population 2 | Dxy | Fst |
| --- | --- | --- | --- |
| Chica Pool1 | Chica Pool2 | 0.0021 | 0.0000 |
| Chica Pool1 | Río Choy | 0.0033 | 0.1440 |
| Chica Pool1 | Pachón | 0.0030 | 0.3501 |
| Chica Pool1 | Rascón | 0.0030 | 0.2592 |
| Chica Pool1 | Los Sabinos | 0.0024 | 0.2029 |
| Chica Pool1 | Tinaja | 0.0024 | 0.2364 |
| Chica Pool2 | Río Choy | 0.0033 | 0.1651 |
| Chica Pool2 | Pachón | 0.0030 | 0.3693 |
| Chica Pool2 | Rascón | 0.0030 | 0.2923 |
| Chica Pool2 | Los Sabinos | 0.0024 | 0.1154 |
| Chica Pool2 | Tinaja | 0.0024 | 0.2477 |
| Río Choy | Pachón | 0.0035 | 0.3654 |
| Río Choy | Rascón | 0.0033 | 0.2507 |
| Río Choy | Sabinos | 0.0034 | 0.1900 |
| Río Choy | Tinaja | 0.0034 | 0.3555 |
| Pachón | Rascón | 0.0026 | 0.4572 |
| Pachón | Los Sabinos | 0.0020 | 0.2832 |
| Pachón | Tinaja | 0.0020 | 0.4430 |
| Rascón | Los Sabinos | 0.0023 | 0.2002 |
| Rascón | Tinaja | 0.0023 | 0.4003 |
| Los Sabinos | Tinaja | 0.0008 | 0.0164 |

**Extended Data Table 6.** Results of D statistic and f4 ratio tests for introgression. *A. aeneus* served as the outgroup. BBAA = derived alleles shared by P1 and P2. ABBA = derived alleles shared by P2 and P3. BABA = derived alleles shared by P1 and P3. Significant p-values (<0.05) indicate evidence of introgression between Choy, Chica, Rascón, and Tinaja.

| P1 | P2 | P3 | D statistic | Z score | p-value | f4 ratio | BBAA | ABBA | BABA |
| --- | --- | --- | --- | --- | --- | --- | --- | --- | --- |
| Río Choy | Chica | Rascón | 0.27893 | 21.4698 | 0 | 0.173087 | 8409.82 | 4100.59 | 2311.94 |
| Río Choy | Tinaja | Chica | 0.14629 | 7.98383 | 7.09E-16 | 0.262394 | 2195.94 | 9381.5 | 6986.95 |
| Chica | Tinaja | Rascón | 0.409914 | 33.6245 | 0 | 0.230212 | 8116.3 | 3375 | 1412.52 |
| Rascón | Tinaja | Río Choy | 0.2558 | 23.5379 | 0 | 0.11872 | 5652.15 | 3207.89 | 1901.03 |

**Extended Data Table 7.** Summary statistics for minor parent (i.e., Río Choy surface fish) ancestry tract lengths in base pair and estimated mean ± SE number of generations since the onset of admixture (T_admix_) in Chica cave Pool 1 (n=5) and Pool 2 (n=14).

| Population | Min | Max | Median | Mean | SE | $T_{\text{admix}}$ |
| --- | --- | --- | --- | --- | --- | --- |
| Pool 1 | 1,350 | 2,069,769 | 29,372 | 48,042 | 263 | 2315±19 |
| Pool 2 | 1,031 | 2,077,917 | 29,561 | 48,435 | 162 | 2294±17 |

**Extended Data Table 8.** The mean minor parent (i.e. surface) tract lengths and mean major parent (i.e. cave) global ancestry proportions were used to infer the timing since admixture (T_admix_).

| Pool | ID | MeanSurface<br>TractLengthBP | MeanSurfaceTract<br>LengthMorgans | MeanCaveGlobal<br>Ancestry | $T_{\text{admix}}$ |
| --- | --- | --- | --- | --- | --- |
| 1 | Chica5_1 | 47096.4 | 0.000546 | 0.770 | 2377.275 |
| 1 | Chica5_2 | 49508.8 | 0.000574 | 0.759 | 2294.262 |
| 1 | Chica5_3 | 49475.3 | 0.000574 | 0.750 | 2324.321 |
| 1 | Chica5_4 | 51214.4 | 0.000594 | 0.744 | 2261.418 |
| 1 | Chica5_5 | 49574.2 | 0.000575 | 0.751 | 2315.212 |
| 2 | Chica1_1 | 51893.7 | 0.000602 | 0.739 | 2246.508 |
| 2 | Chica1_10 | 48084.2 | 0.000558 | 0.755 | 2374.725 |
| 2 | Chica1_11 | 48674.3 | 0.000565 | 0.762 | 2323.384 |
| 2 | Chica1_12 | 47755.7 | 0.000554 | 0.755 | 2389.959 |
| 2 | Chica1_13 | 51812.5 | 0.000601 | 0.768 | 2166.720 |
| 2 | Chica1_14 | 52055.8 | 0.000604 | 0.755 | 2193.243 |
| 2 | Chica1_2 | 50207.8 | 0.000582 | 0.761 | 2257.289 |
| 2 | Chica1_3 | 48756.4 | 0.000566 | 0.754 | 2343.562 |
| 2 | Chica1_4 | 49882.2 | 0.000579 | 0.755 | 2289.383 |
| 2 | Chica1_5 | 48534.3 | 0.000563 | 0.767 | 2315.360 |
| 2 | Chica1_6 | 49014.7 | 0.000569 | 0.773 | 2276.677 |
| 2 | Chica1_7 | 48674.4 | 0.000565 | 0.755 | 2345.607 |
| 2 | Chica1_8 | 49867.9 | 0.000578 | 0.741 | 2333.061 |
| 2 | Chica1_9 | 51125.9 | 0.000593 | 0.745 | 2262.986 |

**Extended Data Table 9.** Absolute genetic divergence (Dxy) for each gene in comparisons between Chica Pool 1 and Pool 2, Choy and Tinaja, Río Choy and Pachón, and Rascón and Pachón.

https://docs.google.com/spreadsheets/d/1yottC4COSed0BGbgjfWui0e4RLgnYa0Hiq_YO_fCc3k/edit#gid=2005687085

**Extended Data Table 10.** Results of GO Analysis on genes with highest genetic divergence (top 5% Dxy) between Chica Pool 1 and Chica Pool 2.

https://drive.google.com/file/d/1RFayjPilvs8D-aXMa28abXl6Ghc5dx5t/view?usp=sharing

**Extended Data Table 11.** Gene descriptions, phenotypes, and results of differential expression, SIFT, VEP, and diploS/HIC selection analyses for candidate genes with ontologies related to cave-adapted phenotypes and in the top 5% of Dxy values between Chica pools. https://docs.google.com/spreadsheets/d/170S6uCIeErV--V5uEzRDpUexXvwB5H_Y59_bD0_hTDk/edit?usp=sharing

**Extended Data Table 12.** Results of differential expression analysis across all time points.

https://docs.google.com/spreadsheets/d/1Xmtlj965TAzRtgz7iQBEDCto132VceVlFK2_er7D3TQ/edit?usp=sharing

**Extended Data Table 13.**
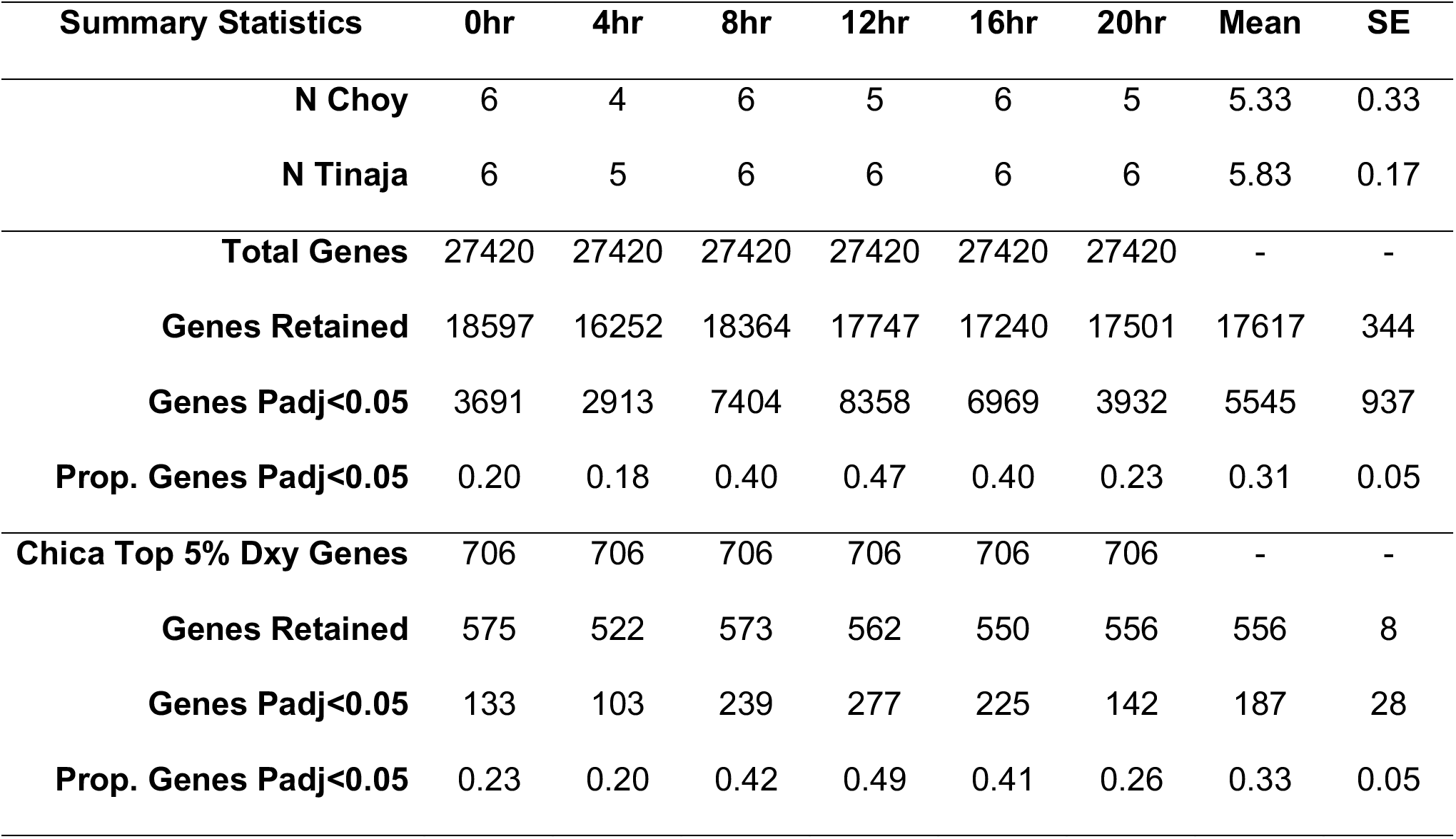
Summary statistics for differential expression analysis in 30 dpf fry from lab-raised cavefish (Tinaja cave) versus surface fish (Río Choy) stock across six timepoints. Summaries are shown for the entire dataset of all genes and a subset of genes within the top 5% of Dxy values between Chica pools.

## Extended Data Figures

**Extended Data Figure 1.**
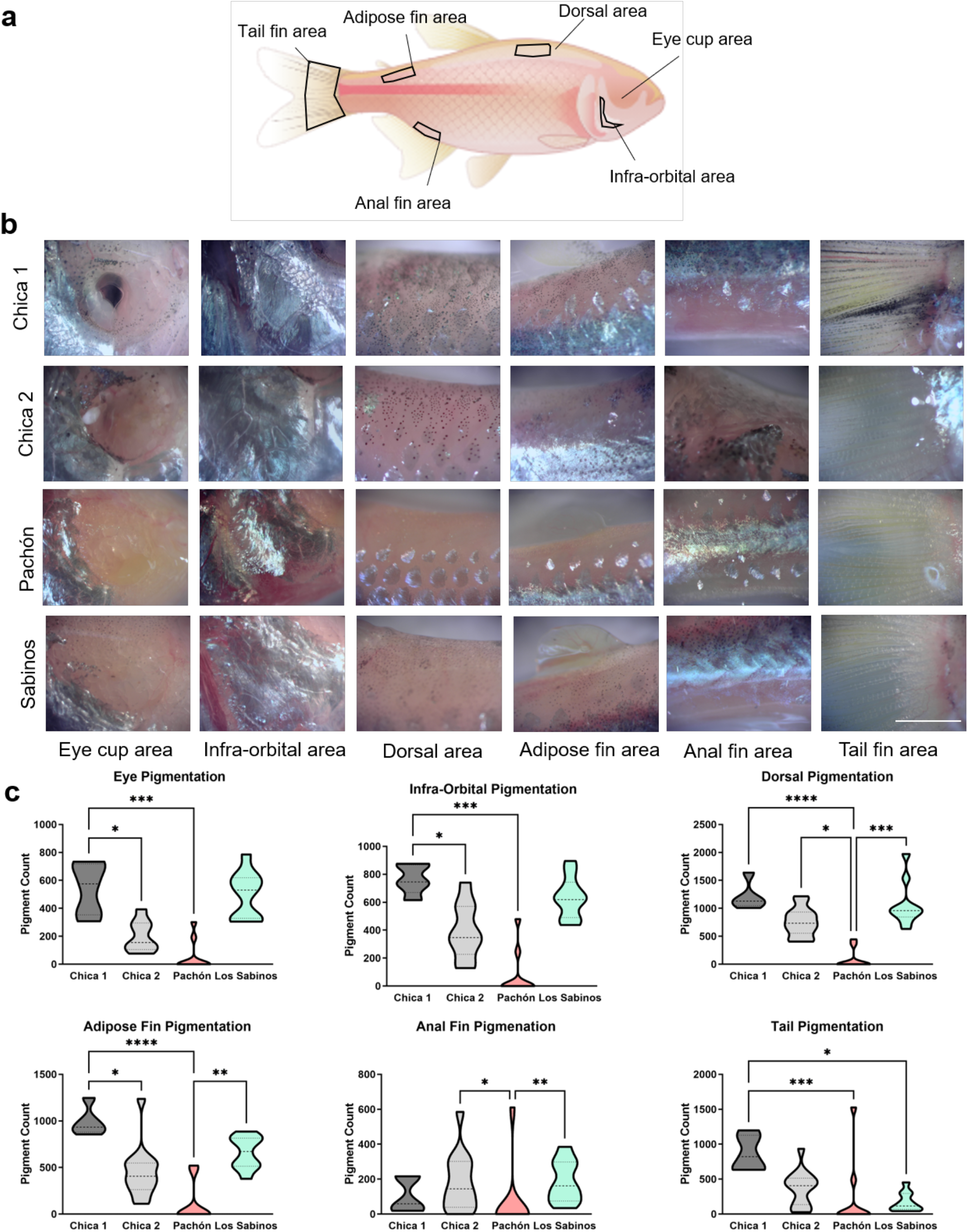
Eye and pigment morphology variations among cave populations. (A) Diagram of areas used for pigmentation quantifications. (B) Brightfield images of fish bodies showing pigmentation across multiple cave populations. (C) Pigment count quantifications by area. Eye cup: Kruskal-Wallis test, P < 0.001, KW statistic = 24.28. Chica 1 vs Chica 2, p<0.05; Chica 1 vs Pachon, p<0.001. Infra-Orbnital: Kruskal-Wallis test, P < 0.001, KW statistic = 22.70. Chica 1 vs Chica 2, p<0.05; Chica 1 vs Pachon, p<0.001. Dorsal area: Kruskal-Wallis test, P < 0.001, KW statistic = 25.66. Chica 1 vs Pachon, p<0.0.001; Chica 2 vs Pachon, p<0.05; Pachon vs Los Sabinos, p<0.001. Adipose area: Kruskal-Wallis test, P < 0.001, KW statistic = 23.65. Chica 1 vs Chica 2, p<0.05; Chica 1 vs Pachon, p<0.001, Pachon vs Los Sabinos, p<0.01. Anal fin: Kruskal-Wallis test, P < 0.01, KW statistic = 12.44. Chica 2 vs Pachon, p<0.05, Pachon vs Los Sabinos, p<0.01. Tail area: Kruskal-Wallis test, P < 0.001, KW statistic = 16.87. Chica 1 vs Pachon, p<0.001, Chica 1 vs Los Sabinos, p<0.05.

**Extended Data Figure 2.**
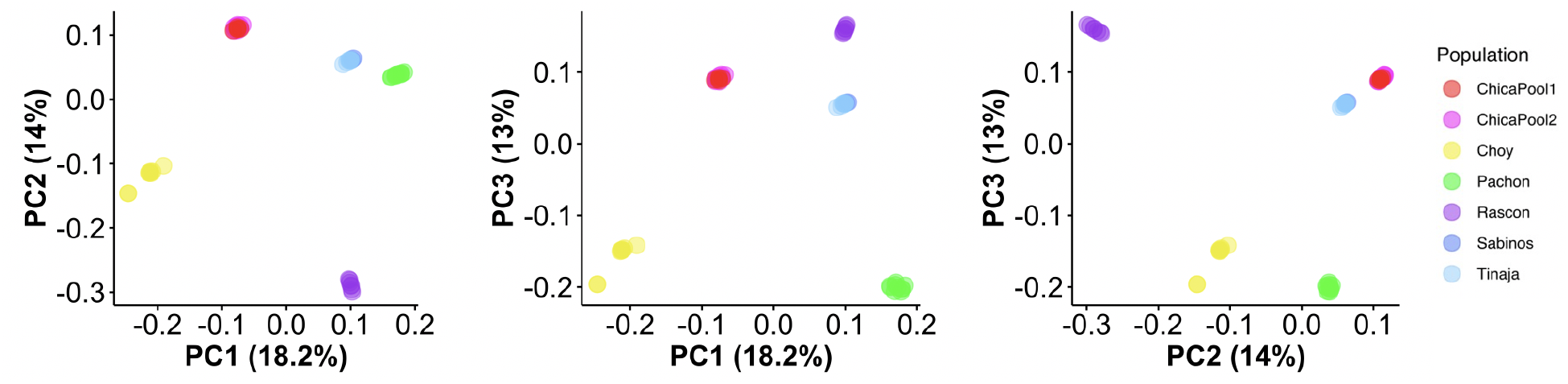
Biplots of scores for the first three PCs from the PCA on SNPs from cave (Chica, Pachón, Tinaja, Los Sabinos) and surface (Río Choy and Rascón) populations. Note that individuals from Chica Pool 1 and Pool 2 overlap and individuals from Tinaja and Los Sabinos overlap.

**Extended Data Figure 3.**
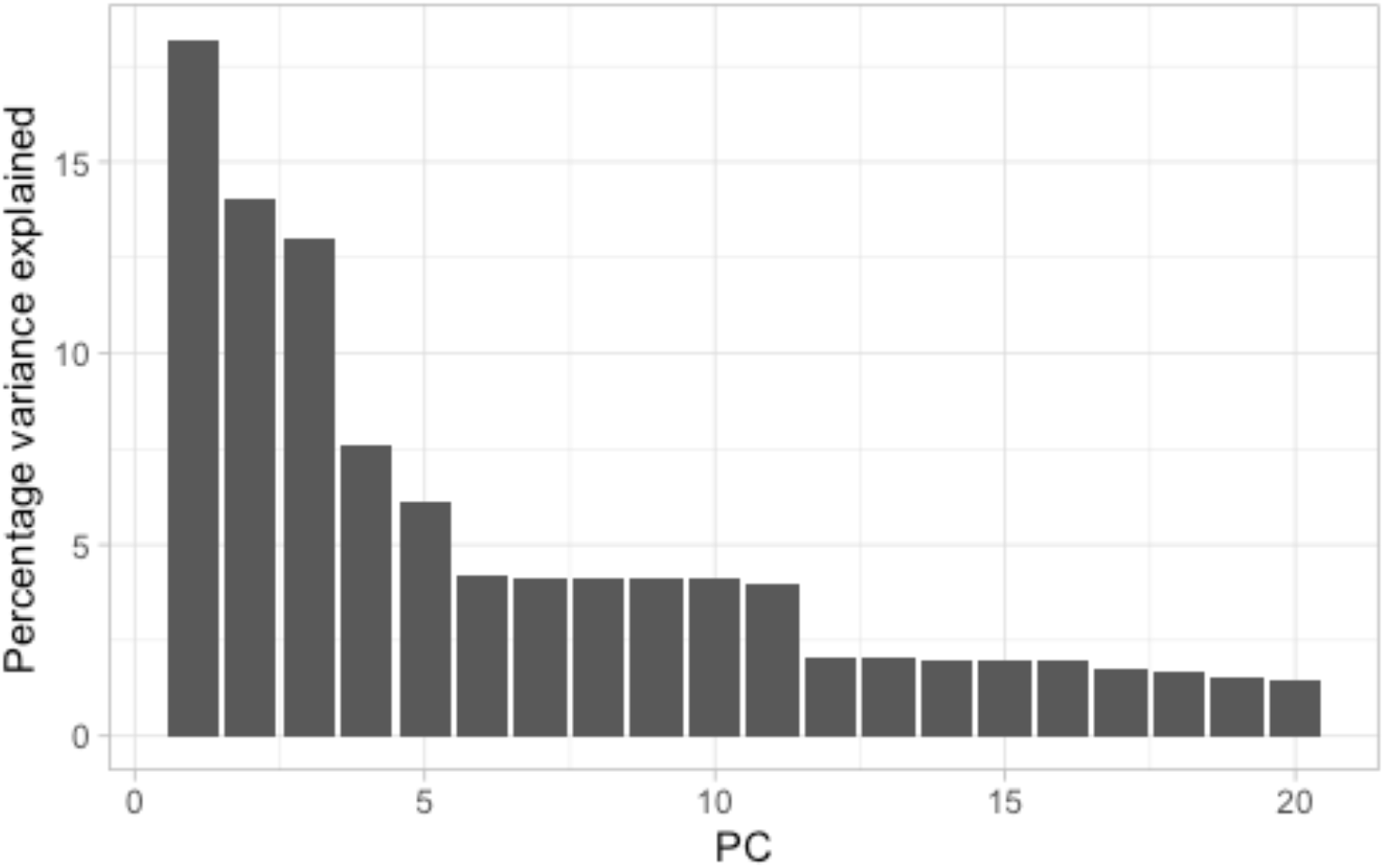
Percentage of variance explained for PCs 1-20 from PCA containing all six *Astyanax* populations (i.e., Chica, Río Choy, Los Sabinos, Pachón, Rascón, and Tinaja).

**Extended Data Figure 4.**
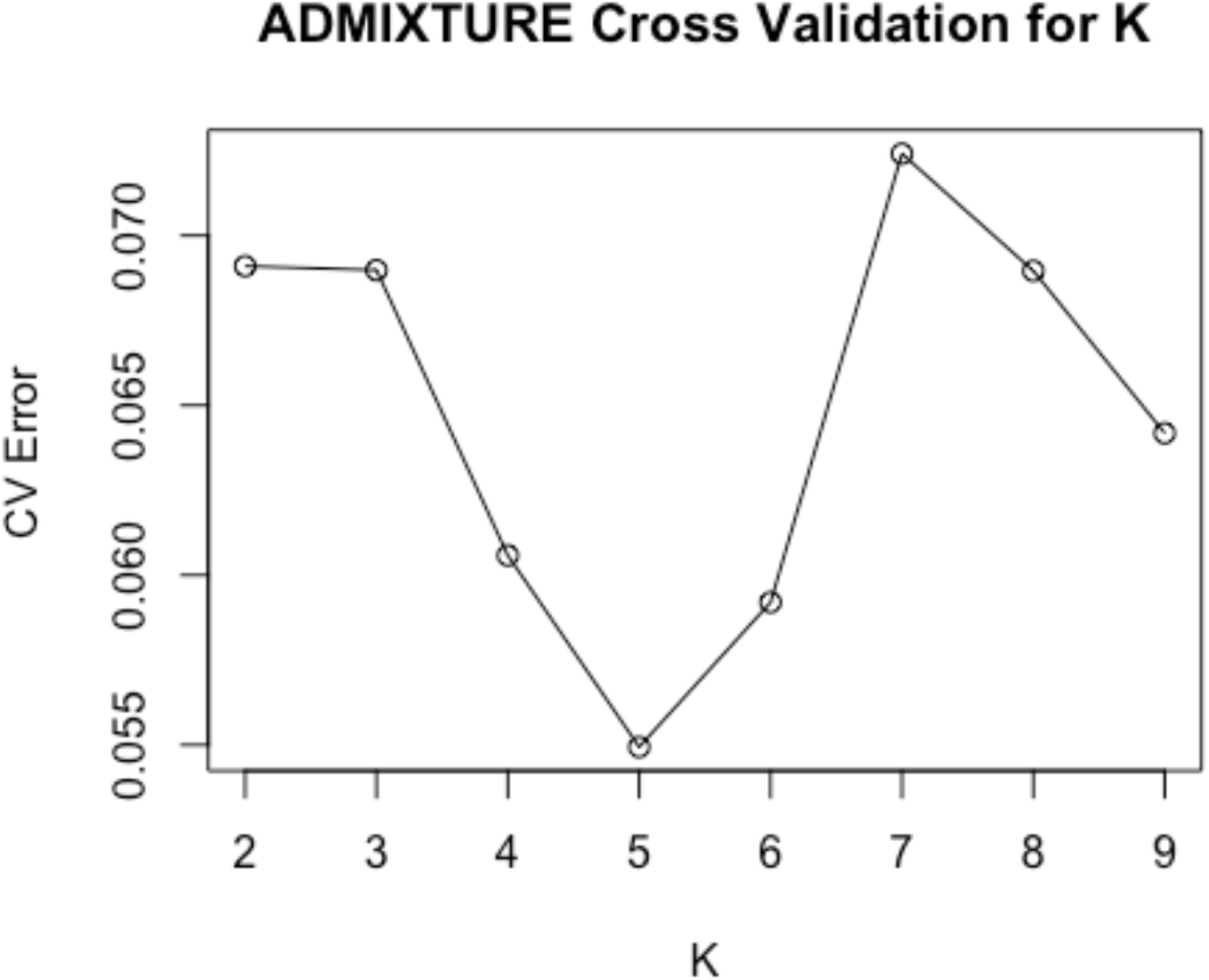
Cross validation (CV) error calculated in ADMIXTURE for values of K ranging from 2-9. A value of 5 is indicated to be the best estimate for the true number of populations clusters because it exhibits the lowest CV error.

**Extended Data Figure 5.**
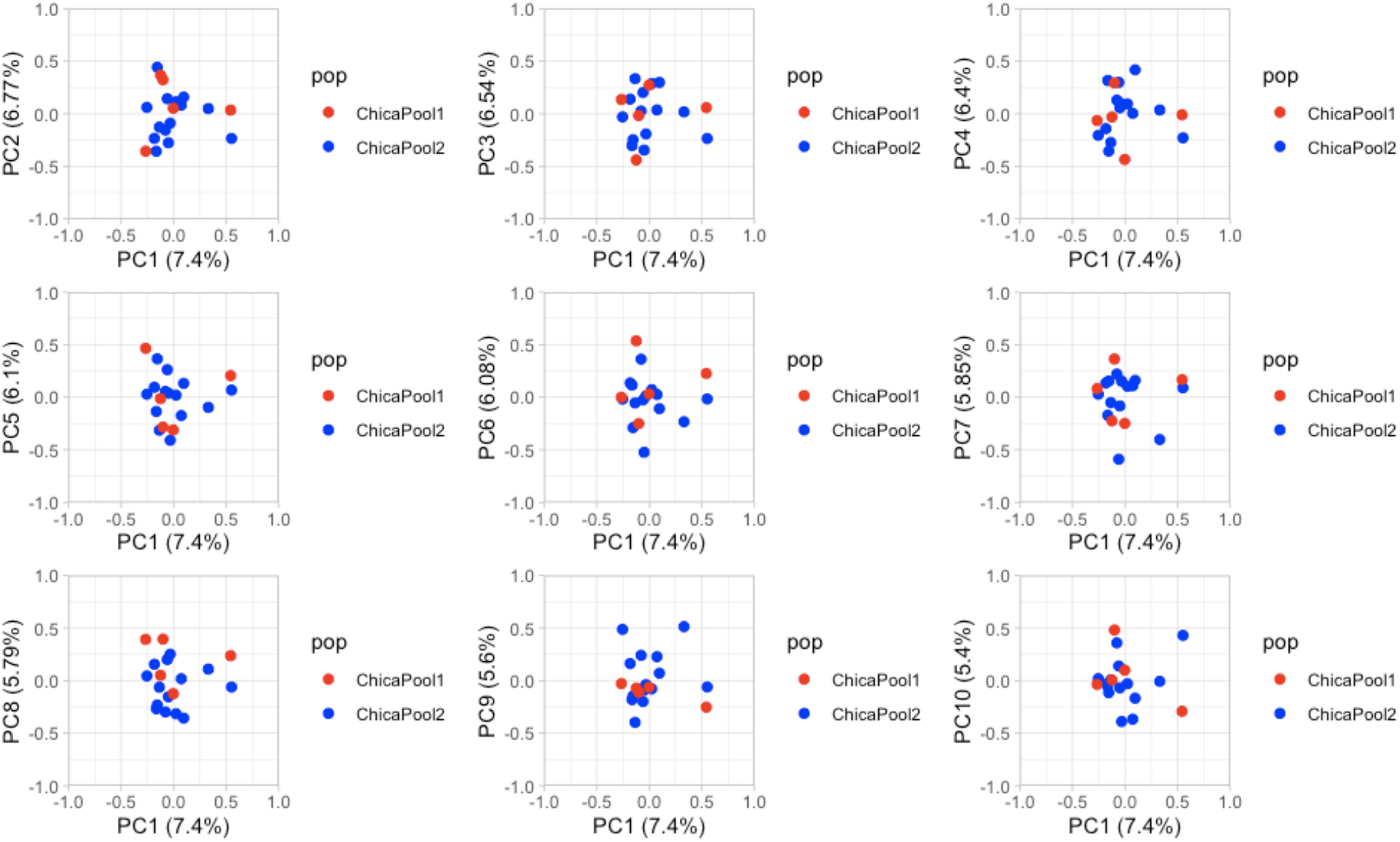
Biplots of PCs 1-10 for PCA including only Chica Pools 1 and 2. Note overlap of individuals from both pools.

**Extended Data Figure 6.**
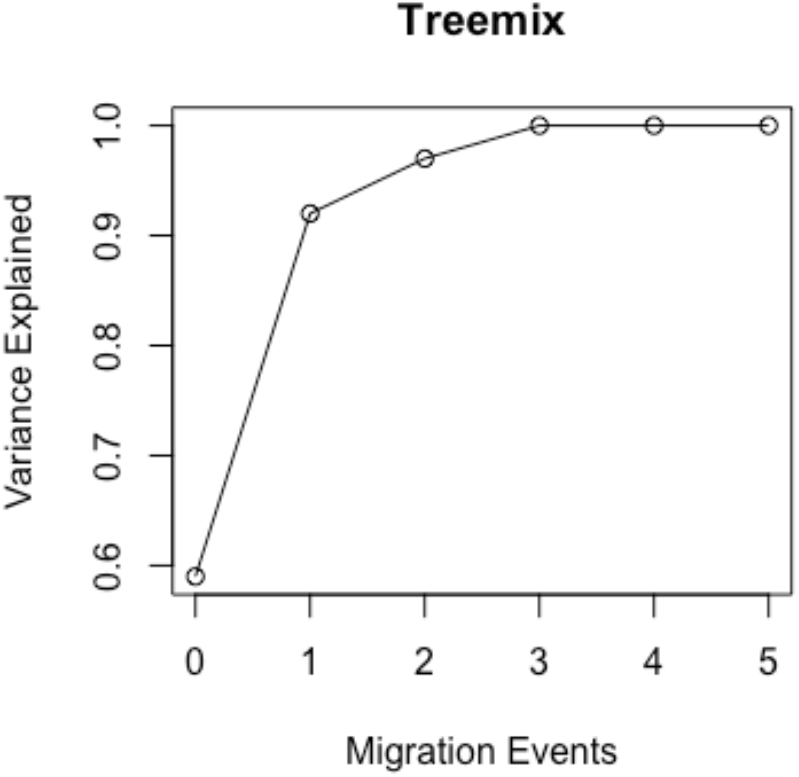
Variance explained for 0-5 migration events in Treemix. Variance explained plateaus at 3 migration events.

**Extended Data Figure 7.**
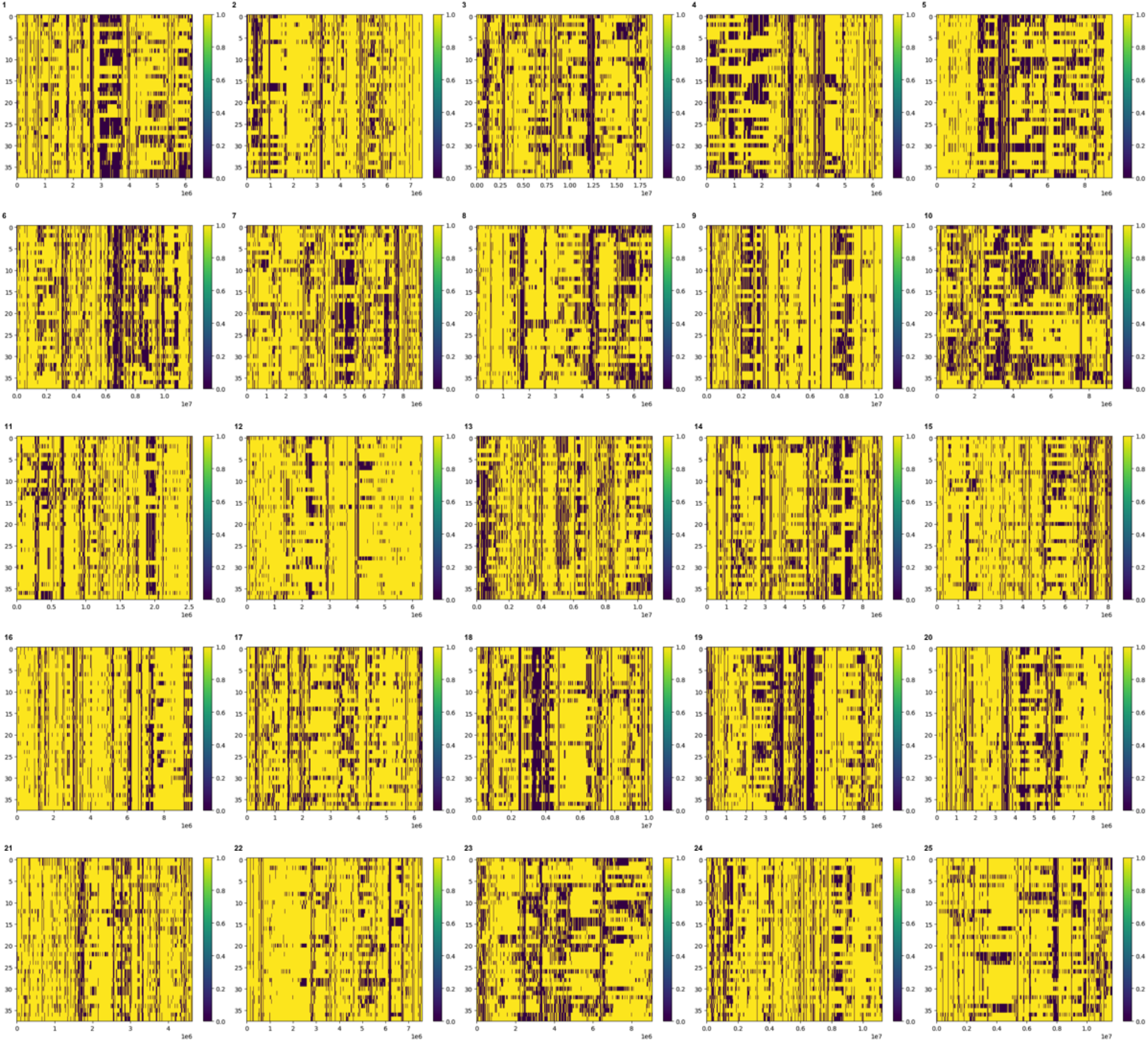
Local ancestry tracts in Chica samples inferred using a Hidden Markov Model approach along each of the 25 chromosomes. Yellow represents cave ancestry and purple represents surface ancestry. The y axis shows haplotypes 1 - 38, with haplotypes 0 - 27 corresponding to Chica Pool 2 (n = 14 diploid individuals), and haplotypes 28 - 38 corresponding to Chica Pool 1 (n = 5 diploid individuals). The x axis shows bp position along each chromosome.

**Extended Data Figure 8.**
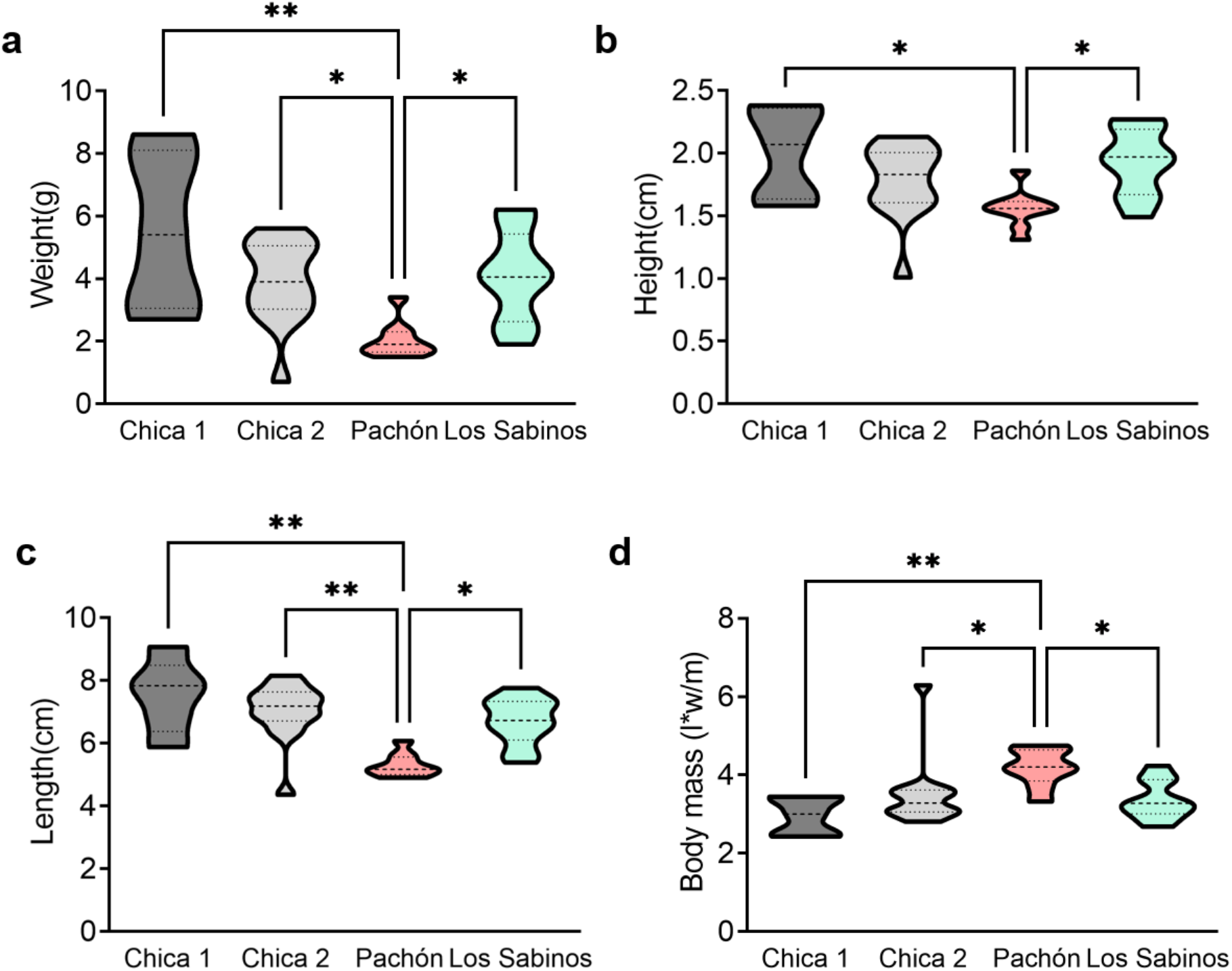
Physical morphology in cave populations of *A. mexicanus*. A. Pachón cavefish weigh significantly less than Chica pool 1 and 2 and Los Sabinos Kruskal-Wallis test, P < 0.01, KW statistic = 15.19. Chica 1 vs Pachon, p<0.01; Chica 2 vs Pachon, p<0.05, Pachon vs Los Sabinos, p<0.05. B. Body height from dorsal fin to stomach is smaller in Pachón cavefish compared to Chica Pool 1 and 2 as well as Los Sabinos. Kruskal-Wallis test, P < 0.01, KW statistic = 11.90. Chica 1 vs Pachon, p<0.05; Pachon vs Los Sabinos, p<0.05 C. Body length measured from mouth to tail is significantly smaller in Pachón cavefish compared to all other cave populations. Kruskal-Wallis test, P < 0.001, KW statistic = 17.84. Chica 1 vs Pachon, p<0.01; Chica 2 vs Pachon, p<0.01, Pachon vs Los Sabinos, p<0.05. D. Body mass is significantly larger in Pachón cavefish compared to other populations Kruskal-Wallis test, P < 0.01, KW statistic *=* 14.41. Chica 1 vs Pachon, p<0.01; Chica 2 vs Pachon, p<0.05, Pachon vs Los Sabinos, p<0.05.

